# Conserved Functional Traits of the Atc Protein System

**DOI:** 10.64898/2026.09.07.749886

**Authors:** Safa Boussouar, Olivier Genest, Mathieu E. Rebeaud, Sébastien Dementin

**Affiliations:** Aix Marseille Univ, CNRS, BIP UMR 7281, IMM, 31 Chemin Joseph Aiguier, 13402 Marseille, France; Institute of Physics, School of Basic Sciences, École Polytechnique Fédérale de Lausanne (EPFL), Lausanne, Switzerland

**Keywords:** Keyword: Molecular chaperone, DnaK, J-domain protein, RNA polymerase, phylogenetic distribution

## Abstract

The Atc protein system (AtcJ, AtcA, AtcB and AtcC) plays a major role in cold adaptation in *Shewanella oneidensis*. AtcJ, a J-domain protein, interacts with the DnaK chaperone and binds AtcC through a conserved C-terminal PX_7_W motif in which Trp is crucial. Overproduced AtcB inhibits RNA polymerase, while the AtcJ-AtcC complex recruits DnaK to modulate this inhibition. This study investigates the conservation of these functional traits across divergent bacteria. We characterized several non-canonical Atc systems and demonstrated that the core interaction network remains functionally preserved despite remarkable variability in the AtcJ C-terminal motif, ranging from intact PX_7_W to degenerate or absent forms, but also in the size of AtcC. Phylogenetic analysis revealed vertical inheritance and co-evolution with host transcriptional machinery. AtcB, AtcC, and AtcJ exhibit strong phylogenetic congruence with the species tree, while AtcA shows greater evolutionary flexibility. The system is enriched in aquatic and psychrophilic lineages but absent in thermophiles. Conditional toxicity assays revealed that AtcB-RNAP interaction depends on specific structural determinants, with DnaK recruitment alleviating toxicity. These findings support a unified molecular mechanism coupling chaperone activity to transcriptional regulation under environmental stress, suggesting that the Atc system represents an ancient and adaptable regulatory module in bacteria.

## Introduction

Protein quality control, and more specifically molecular chaperones, is considered a master modulator of molecular evolution in bacteria, as well as a buffer against deleterious mutations and a key factor in proteome evolution, with the Hsp70 protein often regarded as its central hub (Rebeaud *et al*. 2021; Diaz Arenas *et al*. 2025; Tokuriki and Tawfik 2009; Cowen and Lindquist 2005). The 70 kDa ATP-dependent heat shock proteins (Hsp70s), known as DnaK in bacteria, act as molecular chaperones that play a crucial role in maintaining protein quality control within cells. Their functions range from assisting in the folding of newly synthesized proteins and preventing protein aggregation to dissolving protein aggregates and disassembling protein complexes (Mayer 2021). In *E. coli*, DnaK interacts with over 700 protein substrates, highlighting its central role in protein homeostasis and its essential contribution to cellular survival (Calloni *et al*. 2012). By regulating these processes, DnaK influences a variety of critical cellular functions, including heat shock response, cold adaptation, motility, biofilm formation, and the production of virulence factors, all of which are vital for bacterial adaptation to environmental changes (Mayer 2021; Rebeaud 2026). The activity of DnaK is modulated by J-domain co-chaperone proteins (JDPs), named for their homology to the *E. coli* prototype DnaJ. JDPs are characterized by a conserved J-domain, a ∼70-amino-acid region composed of four helices that form a hairpin structure. The interaction between JDPs and DnaK is mediated by the HPD motif, located in the J-domain (Kampinga and Craig 2010; Craig and Marszalek 2017; Rosenzweig *et al*. 2019). When JDPs bind to DnaK, they induce allosteric changes that stimulate DnaK’s ATPase activity (Kityk, Kopp and Mayer 2018). Beyond regulating DnaK, JDPs act as targeting factors, directing the chaperone toward partially folded, misfolded, or aggregated client proteins. The additional domains in JDPs determine the specificity of these substrate interactions. JDPs also help localize DnaK/Hsp70 to specific cellular compartments where chaperone activity is needed (Mayer and Gierasch 2019; Balchin, Hayer-Hartl and Hartl 2020; Serlidaki *et al*. 2020; Mayer 2026). Recent research suggests that bacteria harbor a vast and unexplored diversity of JDP functions, implying that DnaK may be involved in even more cellular processes than previously recognized (Barriot *et al*. 2020; Malinverni *et al*. 2023).

The *atc* operon of *Shewanella oneidensis* comprises four genes (**Figure 1A**), including *atcJ*, which encodes a short JDP interacting with DnaK through its HPD motif. The functions of *atcA*, *atcB*, and *atcC* remain unknown. Individual deletion of *atcJ*, *atcB*, or *atcC* but not *atcA* results in a growth defect at low temperatures, consistent with the Atc acronym, which stands for adaptation to cold (Maillot *et al*. 2019). AtcJ is composed of the J-domain with a C-terminal extension of 21 amino-acids which is necessary to strongly interact with AtcC (Kd in the order of 10 nM). The tryptophan of the conserved PX_7_W motif in this C-terminal peptide is essential for binding with AtcC (Weber *et al*. 2023). AtcC interacts with AtcB, but the molecular determinants of this interaction remain unknown. Interestingly, AtcB binds RNA polymerase (RNAP) through two conserved Asp residues, most probably at the level of a positively charged alpha helix of the RpoC subunit of the polymerase (Boussouar *et al*. 2025). This interaction is essential for the function of the operon since a *S. oneidensis* strain carrying the chromosomal copy of a variant of AtcB in which the two Asp have been replaced by Ala is unable to grow at low temperature. Nevertheless, under conditions of overproduction (in *S. oneidensis* or *E. coli*), AtcB is toxic because it stably binds to RNA polymerase (RNAP), thereby preventing cellular transcription. However, the amount of inactive AtcB-RNAP complex considerably decreases in the presence of the partners AtcC and AtcJ, which, in complex, recruit DnaK (**Figure 1B**). This observation has led us to propose that the Atc protein system couples the chaperone activity of DnaK to transcription to regulate specific genes involved in cold growth (Maillot *et al*. 2021; Boussouar *et al*. 2025). However, the underlying mechanism and the targeted genes remain to be discovered.

**Figure 1:**
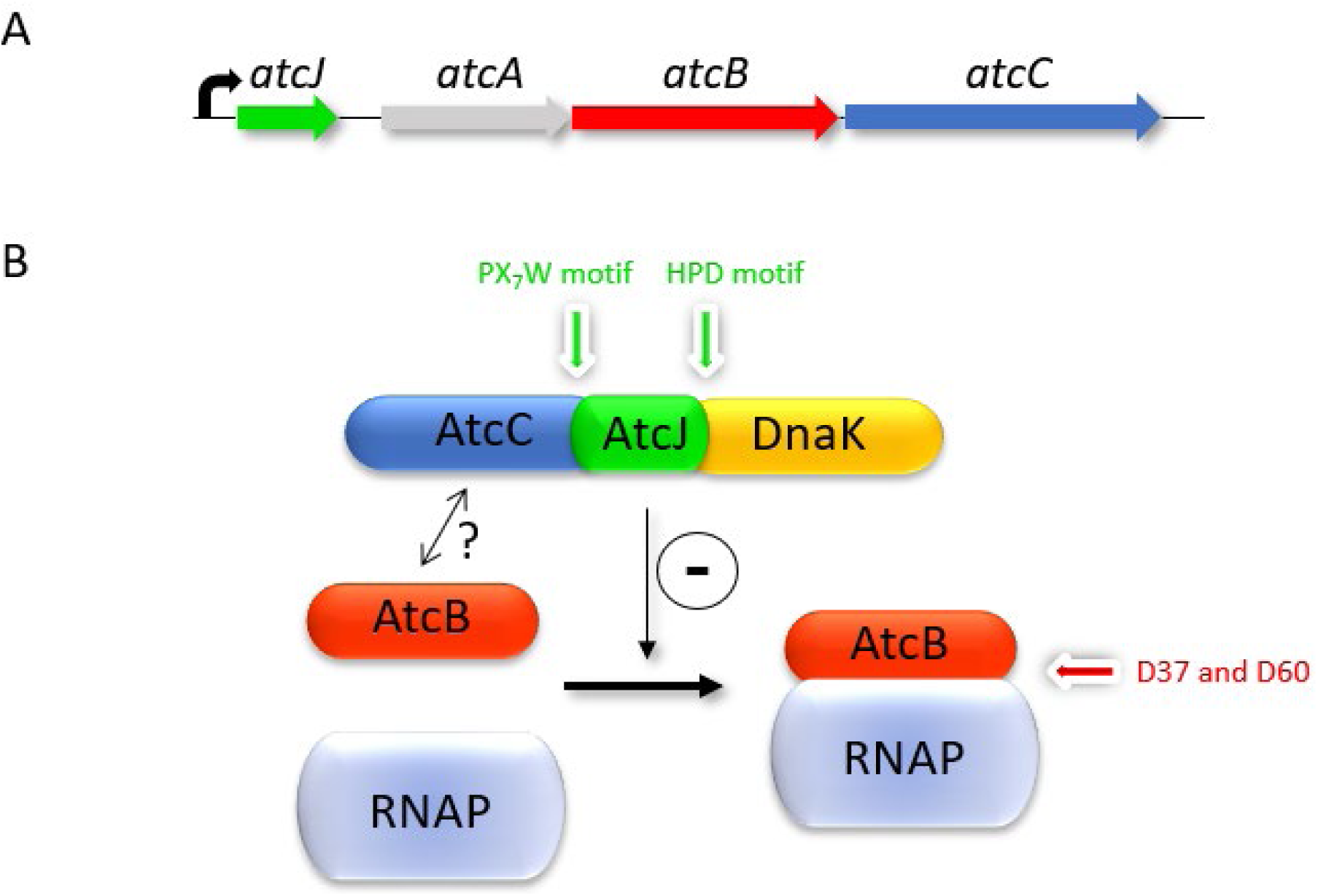
The *atc* operon from *S. oneidensis*. (A) *atc* operon. (B) Functional protein interaction network. DnaK is recruited by AtcJ through the HPD motif of the J-domain. The conserved W89 of the 21 amino acids C-terminal extension is essential for strong interaction with AtcC which also interacts with AtcB. Under overproduction conditions, AtcB interacts with and inhibits RNAP through the two conserved D37 and D60 residues, but the amount of AtcB-RNAP complex is limited by DnaK which is recruited by the AtcJ-AtcC complex. Under physiological conditions, the AtcJ-DnaK, AtcJ-AtcC and AtcB-RNAP interactions are crucial for growth at low temperature.

A BLAST-P analysis to identify homologs of the Atc proteins led to the observation that the *atc* operon is present in organisms other than *S. oneidensis,* mainly in environmental Gamma- and Beta-proteobacteria, especially aquatic ones. Interestingly, the four genes are always in the same order. Notable differences are observed regarding the length of the *atcC* gene and the possible presence of additional genes in the operon. Additionally, in some operons, the canonical PX_7_W sequence of the C-terminal peptide of AtcJ is divergent (lacking Pro and/or Trp) or absent (Weber *et al*. 2023).

Thus, we questioned whether a conserved mechanism underlies the Atc system by comparing the properties of divergent Atc systems relative to the canonical system of *S. oneidensis*. Additionally, by conducting a formal phylogenetic study, we sought to determine the distribution of the Atc system across the bacterial world. Our data show that the interaction network verifies the rules determined with the *Shewanella* system. Interestingly, not all AtcB proteins bound *E. coli* RNAP. This was rationalized by our phylogenetic analysis, which shows that the specificity of an AtcB protein towards RNAP in a particular organism is probably dictated by specific molecular and structural features constrained by RNAP.

## Methods

### Strains, plasmids and primers

Strains of *E. coli*, plasmids and primers used in this study are described in **Supplementary Material**. Transformations of *E. coli* strains were performed using the heat shock method, with cells made competent by cold CaCl_2_ treatment.

### Source of DNA

The genes coding for the AtcJ, AtcB and AtcC proteins from *Collimonas arenae* Cal35 (Care) were amplified from genomic DNA kindly provided by Dr Stéphane Uroz (INRAE, Nancy, France). Genomic DNA from *Solidesulfovibrio magneticus* RS-1 (Dma) was extracted from cells of a culture kindly provided by Dr Christopher Lefèvre (Biosciences and Biotechnologies Institute (BIAM), CEA, Cadarache, France), using Monarch Genomic DNA Purification Kit (New England Biolabs) and was used as template to amplify the *atcJ*, *atcB* and *atcC* genes. The synthetic *atcJ*, *atcB* and *atcC* genes from *Janthinobacterium* sp. B9-8 (Jab) optimized for protein production in *E. coli* were obtained from IDT (Integrated DNA technology). The synthetic *atcB* genes from *Thiodictyon syntrophicum* Cad16T (Tsy) and *Verrucomicrobium* sp. GAS474 (Veg) optimized for protein production in *E. coli* were also obtained from IDT (Integrated DNA technology). The primers and strategy used for cloning genes into pBad33, pBad24-CBP, pT18 and pT25 vectors are described in **Supplementary Material**.

### Growth conditions

Strains were cultured in Luria Bertani (LB) rich medium, comprising bacto-tryptone (10 g/L), yeast extract (5 g/L), and NaCl (5 g/L) at pH 7.4. When antibiotic selection was required, the following concentrations were used: chloramphenicol (25 μg/mL) and ampicillin (50 μg/mL). AtcB toxicity assays were run by monitoring growth of *E. coli* MG1655 containing pBad33 plasmids expressing *atcB*, either WT or mutated, in the presence or not of *atcC* and *atcJ* (WT or mutated). Overnight cultures at 28°C in LB medium in the presence of 0.05% glucose were inoculated into 1 mL of fresh medium containing 0.5% arabinose and chloramphenicol to initial optical density at 600 nm (OD_600_) = 0.01 in 24-wells plates. Growth at 28°C under agitation was monitored by measuring OD_600_ using a microplate reader (SPARK 10M, TECAN) over 15h.

### Bacterial two hybrid (BACTH) assays

To assess the *Shewanella oneidensis*, *Collimonas arenae* Cal35 and *Janthinobacterium* sp. B9-8 AtcJ-AtcC and AtcB-AtcC interactions, *E. coli* strain BTH101 was co-transformed with the plasmid constructs pT18-*atcJ* (WT or mutated) or pT18-*atcB* and pT25-*atcC* (**Supplementary material**). Transformants were selected on LB-agar medium containing ampicillin and kanamycin after 48 h (or 72 h for *Janthinobacterium* sp. B9-8 AtcB-AtcC interaction) of incubation at 28°C. The rest of the procedure is as described before (Weber *et al*. 2023).

### Pull-down assays

Interactions of WT or mutated AtcB from *Shewanella oneidensis*, *Collimonas arenae* Cal35, *Janthinobacterium* sp. B9-8, *Thiodictyon syntrophicum* Cad16T, *Verrucomicrobium* sp. GAS474 and *Solidesulfovibrio magneticus* RS-1 with *E. coli* RNAP subunits were assessed by copurification assays. MG1655 or RLG3982 (RpoC_Δ215-220) *E. coli* strains were transformed with pBad24 plasmid coding for calmodulin binding protein (CBP) tagged AtcB from each strain (**Supplementary Material**). To assess the interactions between AtcB and AtcC from *Shewanella oneidensis*, *Collimonas arenae* Cal35, *Janthinobacterium* sp. B9-8 and *Solidesulfovibrio magneticus* RS-1, the *E. coli* strains were co-transformed with pBad24 plasmid coding for CBP-tagged AtcB and pBad33 plasmid coding for AtcC-6His (**Supplementary Material**). The assay was performed with calmodulin beads as previously described (Boussouar *et al*. 2025).

### Western blots

To detect AtcC in its 6His version from pBad33-derived plasmids in pull-down fractions, the samples from the same amount of cells were heat-denatured, loaded on SDS-PAGE, transferred by western blot and revealed with an anti-His antibody (6x-His Tag Monoclonal Antibody (AD1.1.10), HRP, # MA1-80218, ThermoFisher Scientific).

### Phylogenetic analysis and search for Atc-containing organisms

Homologs of the *atc* operon proteins were identified mostly in Beta- and Gammaproteobacteria by sequence similarity searches against publicly available reference proteomes in Uniprot (UniProt 2025). A species phylogeny of the 294 strains found (**Supplementary File**) was reconstructed from concatenated conserved genomic markers using maximum-likelihood inference with IQ-TREE3, based on the 20 validated core genes (Tian and Imanian 2023) without *rpoA* and *rpoB* (*cgtA, EngA, Ffh, infB, lepA, nusA, pheS, prfA, rplA, rplB, rplC, rplE, rpsB, rpsC, rpsE, rpsG, secY* and *ychF*). A local database was constructed with each proteome of the species harboring the *atc* operon, and a local Blast was performed to extract each protein of interest (**Supplementary File**). Phylogenetic analyses for each operon component (AtcA, AtcB, AtcC, and AtcJ) (**Supplementary File**) were performed using IQ-TREE 3.1.0 (Wong *et al*. 2026). Protein sequences were aligned with MUSCLE3 algorithm (Edgar 2004), and poorly aligned regions were trimmed with trimAl (v1.4) using the -gt (gapthreshold) and -cons (Minimum percentage of positions in the original alignment to conserve) methods (Capella-Gutiérrez, Silla-Martínez and Gabaldón 2009). The best-fitting substitution model was determined by ModelFinder (Kalyaanamoorthy *et al*. 2017) as LG+F+I+G4, which was subsequently used for maximum likelihood tree reconstruction. Branch support was estimated using 1,000 ultrafast bootstrap replicates (Hoang *et al*. 2018) and approximate likelihood-ratio test -aLRT tests (Guindon *et al*. 2010). The resulting trees were visualized and annotated in iTOL (Letunic and Bork 2024).

### Phylogenetic congruence and patristic distances analyses

All Mantel tests and tanglegrams analyses were performed in R (version ≥4.5). Congruence between operon-protein phylogenies and the species phylogeny was assessed visually using tanglegrams and quantitatively by Mantel tests comparing pairwise phylogenetic distance matrices. The patristic (co-phenetic) distances were calculated from the species tree and each operon-protein tree. The distance matrices were restricted to taxa common to the species tree and each protein tree and reorganized to have identical end labels. The pairwise distances were extracted from each matrix and the relationship between protein evolutionary distances and species phylogenetic distances was visualized using linear regression, and the association was determined by R². Packages used were *ape*, *phytools*, *dendextend*, *viridis*, *dplyr*, *phylogram*, and *vegan*.

## Results

### Selection of non canonical Atc systems

Based on a previous P-BLAST analysis (Weber *et al*. 2023), we identified AtcJ homologs that diverge from *Shewanella oneidensis* AtcJ, specifically in that the conserved PX_7_W motif, present in most homologs, is either degenerate or absent. For example, in *Janthinobacterium* sp. B9-8 (Xu *et al*. 2019), *Iodobacter ciconiae* H11R3 (Lee *et al*. 2019), *Iodobacter fluviatilis* PCH194 (Kumar *et al*. 2022), *Deefgea piscis* D17 (Gim *et al*. 2022), and *Dechloromonas denitrificans* G6 (Horn *et al*. 2005), the Trp residue is replaced by Ala, Ser, or Thr. In *Thiodictyon syntrophicum* Cad16T (Luedin *et al*. 2018) and *Massilia* spp., the Pro residue is replaced by Ala. More drastically, the AtcJ proteins from *Verrucomicrobium* sp. GAS474 (Pold *et al*. 2018) and *Solidesulfovibrio magneticus* RS-1 (Shimoshige *et al*. 2021) lack this motif entirely. Nevertheless, these AtcJ proteins are encoded in potential operons alongside the other three Atc proteins (AtcA, AtcB, and AtcC), which share homology with their *S. oneidensis* counterparts. These operons exhibit a genetic organization similar to that of *S. oneidensis*, with the four genes arranged as *atcJ*, *atcA*, *atcB*, and *atcC*. Another interesting aspect is the notable size variation among AtcC homologs, which is not observed with the AtcJ, AtcA and AtcB proteins whose size is more homogeneous. For example, AtcC from *Shewanella oneidensis* which contains 298 amino acids is considerably shorter than other homologs (Weber *et al*. 2023). On these basis, we selected five Atc systems :*Collimonas arenae* Cal35 (Care) (Wu *et al*. 2015), *Janthinobacterium* sp. B9-8 (Jab), *Thiodictyon syntrophicum* Cad16T (Tsy), *Verrucomicrobium* sp. GAS474 (Veg), and *Solidesulfovibrio magneticus* RS-1 (Dma), to characterize their interaction networks and compare them to that of *S. oneidensis*. The features of the selected systems are summarized in **Table S1** and in **Figure 2**.

**Figure 2:**
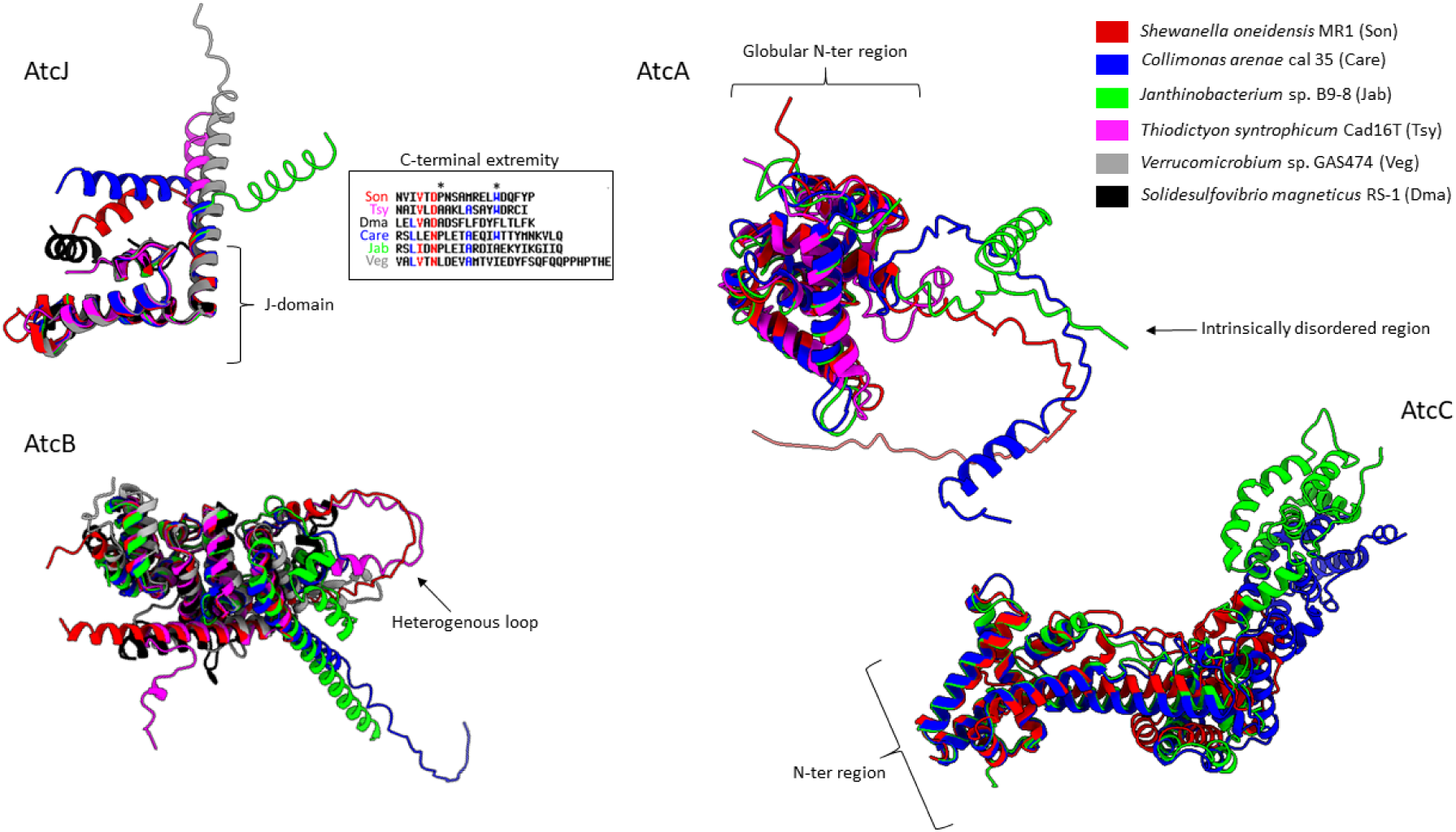
AlphaFold 3 predictions and overlay of the structures of the AtcA, AtcB, and AtcC homologues from the selected organisms. The rank 1 structure models are shown. The sequence of C-terminal extremity of the AtcJ homologues is shown in the box. Model representation and superposition are performed using ChimeraX software. Only the models with pTM >0.5 are shown.

Structural models of the Atc proteins were generated using AlphaFold3, with predicted template modeling (pTM) scores provided in **Table S1**. The AtcJ proteins exhibit a broadly similar predicted structure at the level of their J-domain (**Figure 2**). Their C-terminal peptides are partially folded as an α-helix, although with low confidence for AtcJ from Veg. They contain the PX_7_W motif, either intact (Care) or degenerate (Jab, Tsy), or lack it entirely (Veg, Dma) (**Figure 2**). With the exception of the Veg system, the AtcA proteins contain a predicted C-terminal intrinsically disordered region (IDR) enriched in Asp and Glu residues. Where the confidence is reasonable (pTM > 0.5), the folding of the globular N-terminal region is conserved (**Figure 2**). Regarding the AtcB proteins, all retain the two conserved Asp residues critical for RNAP binding (Boussouar *et al*. 2025) (**Table S1**), and their overall folding remains similar, with the exception of a loop that varies in length depending on the protein (**Figure 2**). The AtcC proteins in the selected Atc systems are substantially longer than that of *S. oneidensis* (**Figure 2** and **Table S1)**). However, where the confidence is sufficient (pTM > 0.5 for AtcC from Son, Care, and Jab), the N-terminal regions exhibit similar folding while their C-terminal regions seem different. This region has been proposed as the interaction site with AtcJ (Weber *et al*. 2023).

### Production of AtcBs in *E. coli*

We initially expressed the selected AtcB proteins in *E. coli* to evaluate (1) their ability to bind RNAP by co-purification experiments and (2) their associated cellular toxicity. For co-purification experiments, as previously described (Maillot *et al*. 2021; Boussouar *et al*. 2025), the proteins were overexpressed as CBP-tagged fusions. Cell extracts were then incubated with calmodulin resin to immobilize AtcB. After washings, AtcB and its interacting partners were eluted and analyzed by SDS-PAGE (**Figure 3A**). Surprisingly, the resin-immobilized CBP-AtcB fusions exhibit migration profiles that are inconsistent with their theoretical sizes. For example, the CBP-AtcB_Care_ fusion (theoretical MW = 29.501 kDa) displays an apparent molecular weight on the gel higher than that of the CBP-AtcB_Jab_, whose theoretical MW is 32.481 kDa. This discrepancy warrants further investigation. As previously reported, RpoA, RpoB, and RpoC co-eluted with AtcB from Son (Maillot *et al*. 2021; Boussouar *et al*. 2025). A similar interaction pattern was observed for AtcB from Care and Jab, but not for AtcB from Tsy, Veg, or Dma. Consistently, overexpression of AtcB from Son, Care, and Jab impaired *E. coli* growth, with the toxicity severity correlating with the amount of co-immobilized RNAP (**Figure 3B**). Because not all AtcB proteins bind *E. coli* RNAP with sufficient affinity to form a stable complex, we hypothesized that the interaction between AtcB and RNAP depends on structural features unique to each specific AtcB-RNAP pair. This will be discussed later in the manuscript.

**Figure 3:**
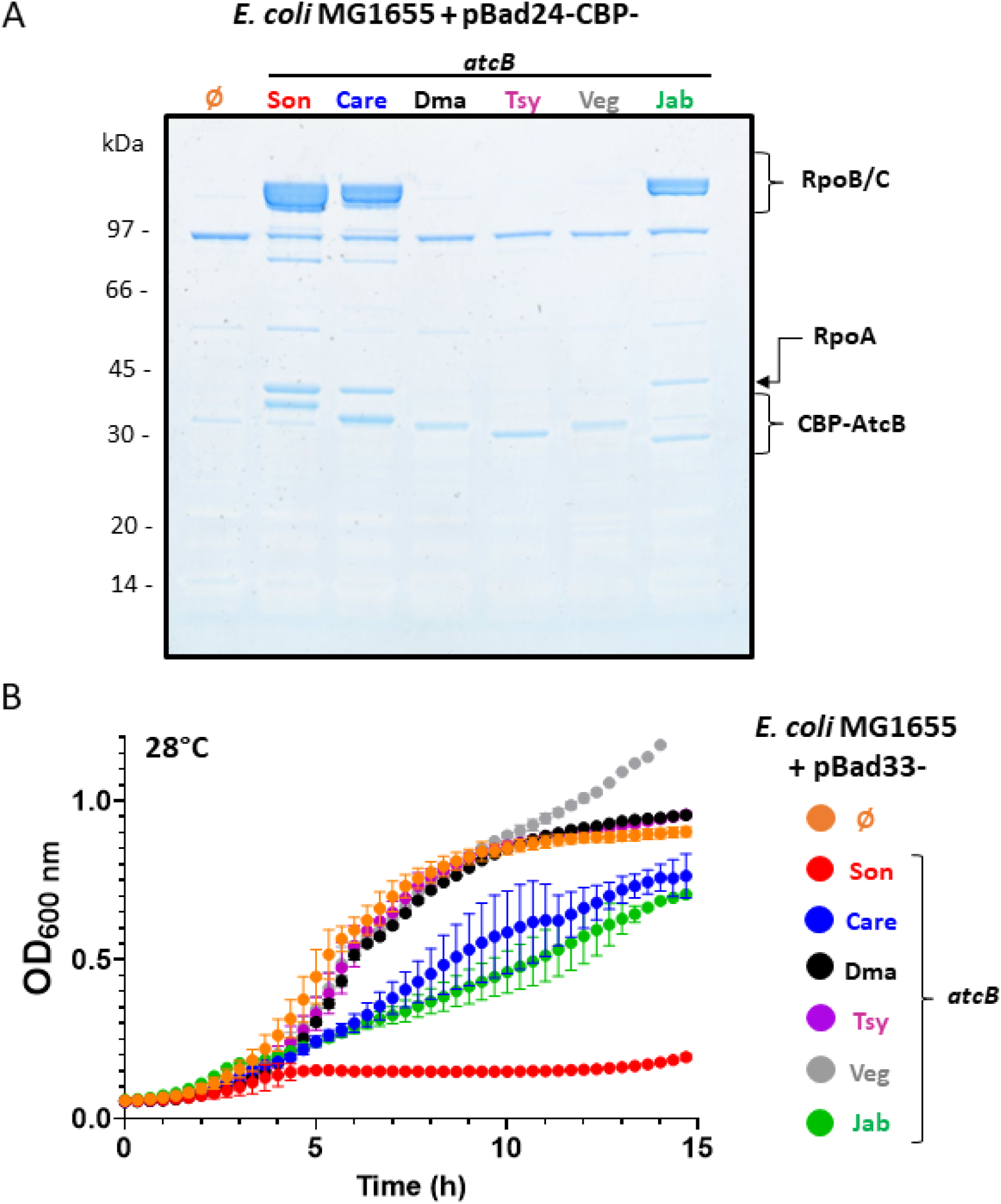
Production of exogenous AtcB proteins in *E. coli*. (A) In vivo co-purification of AtcB proteins and RNAP in *E. coli* MG1655. Strains containing the pBad24 vector allowing the overproduction of CBP-tagged AtcB from *Shewanella oneidensis* (Son)*, Collimonas arenae* Cal35 (Care), *Solidesulfovibrio magneticus* RS1 (Dma), *Thiodictyon syntrophicum* Cad16T (Tsy), *Verrucomicrobium* sp. GAS474 (Veg) and *Janthinobacterium* sp. B9-8 (Jab) were grown at 37°C to the exponential growth phase before 0.2% arabinose was added for 2h. Control experiments with CBP produced alone were included (∅=empty vector). CBP-AtcB was purified on calmodulin beads. CBP-AtcB and the co-purified proteins were separated by SDS-PAGE, and stained with Coomassie blue. (B) Growth of *E. coli* MG1655 at 28°C producing AtcB from the six strains from pBad33. Cultures were conducted in LB medium in the presence of chloramphenicol and 0.5% arabinose. After initial overnight growth at 28°C, strains were diluted to OD_600_ = 0.01, and were incubated at 28°C with shaking. Absorbance was measured over time. The data from two replicates are shown as mean ± SD.

We therefore focused on the two AtcB proteins (from Care and Jab) that bound *E. coli* RNAP. We had previously shown that the AtcB_Son_-RNAP interaction is mediated by the two conserved aspartate residues of AtcB, which interact with a positively charged helix in the RpoC subunit of RNAP (Boussouar *et al*. 2025). We examined this with AtcB proteins from Care and Jab, repeating the co-immobilization and growth experiments described in **Figure 3A** and **3B** using AtcB variants with mutations at these aspartate residues (D28A/D51A variant for AtcB_Care_ and D33A/D56A for AtcB_Jab_). As expected, these variants were unable to bind RNAP (**Figure 4A**) and showed no toxicity (**Figure 4C, D and E)**. Conversely, when we performed co-immobilization experiments in the *E. coli* strain RLG92 (Bartlett *et al*. 1998, 2000), which lacks the positively charged helix in RpoC, WT AtcB from Care and Jab proteins failed to interact with RNAP, confirming that the interaction is also specific to this helix (**Figure 4B**). We also demonstrated that the toxicity of AtcB from *S. oneidensis* is alleviated by DnaK, which is recruited by the AtcJ-AtcC complex (**Figure 4C**) (Boussouar *et al*. 2025). To assess whether this also applied to the Care and Jab Atc systems, we conducted toxicity assays in which AtcB proteins were co-expressed with their AtcC and AtcJ partners. As expected, no AtcB toxicity was observed under these conditions (**Figure 4D** and **4E**). We then repeated these experiments using AtcJ variants: one carrying the H31Q mutation in the HPD motif (unable to recruit DnaK) and another lacking the C-terminal extension (ΔCter mutant, unable to interact with AtcC). In both cases, AtcB toxicity persisted, although partially in the case of Jab, further confirming that DnaK recruitment by the AtcJ-AtcC complex is required to mitigate AtcB toxicity (**Figure 4C, 4D** and **4E**). The C-terminal peptide of AtcJ is essential to mediate the interaction with AtcC, as previously described for the *S. oneidensis* system (Maillot *et al*. 2019; Boussouar *et al*. 2025) and suggested in the predicted AtcJ-AtcC complex structures (**Figures S1A** to **S1F**). To further investigate the molecular determinants of the AtcJ-AtcC interaction, we repeated the toxicity assays using point mutations in residues of the C-terminal peptide of AtcJ that are predicted to mediate the interaction. For AtcJ from *S. oneidensis*, we tested the W89R variant, as the hydrophobic Trp (part of the PX_7_W motif) was previously shown to be essential for the interaction (a structural model of the AtcJ-AtcC complex is shown **Figure S1A**). AtcB toxicity was partially restored in this condition, indeed growth was not totally impaired but strongly retarded (**Figure 4C**). We estimated that this partial effect results from a residual AtcJ-AtcC interaction supported by protein overexpression. For AtcJ from Care, we designed the corresponding W83R variant (see the structural model of the AtcJ-AtcC complex shown in **Figure S1B)**, and obtained this time a complete restoration of AtcB toxicity (**Figure 4D**). To confirm this finding, we used bacterial two-hybrid assay (BACTH), which relies on the reconstitution of the T18 and T25 catalytic domains of *Bordetella pertussis* adenylate cyclase. We constructed two separate plasmids: one fusing AtcJ to the T18 domain and another fusing AtcC to the T25 domain. When both fusion proteins are co-expressed in an *E. coli* strain lacking adenylate cyclase, the interaction between AtcJ and AtcC from Care restores adenylate cyclase activity, leading to cAMP production. This, in turn, activates the *lac* operon, resulting in high β-galactosidase activity, a measurable indicator of the protein-protein interaction. As shown in **Figure S2**, high β-galactosidase activity was measured for WT AtcJ and AtcC proteins, confirming interaction. As observed for the Son Atc system (Weber *et al*. 2023), absence of the C-terminal extension (ΔCter mutant) or replacement of the conserved Trp by Arg (W83R mutation) abolished interaction. Moreover, the experiment performed with the AtcJ C-terminal extension alone fused to T18 confirmed this peptide determined interaction with AtcC. For AtcJ from Jab, lacking the Trp residue which is replaced by Ala (A83), we tested whether this Ala and neighboring hydrophobic residues (I82 and Y86) could compensate for the absence of Trp, as suggested by the structural model shown in **Figure S1C**. We tested the effect of the I82R, A83R and Y86R AtcJ mutations on the restoration of AtcB toxicity in the presence of AtcC (**Figure S3**). The I82R and A83R mutations partially restored toxicity to an extent similar to the H31Q and ΔCter mutations (**Figure 4E**), suggesting that both the I82 and A83 residues determine binding to AtcC. Mutating the tyrosine at position 86 had however no effect, suggesting this residue is not involved in interaction with AtcC. We conducted BACTH studies to support these results but the experiments run with the WT AtcJ and AtcC proteins did not allow us to measure β-galactosidase activity. We assumed that one or both fusion proteins might not be stable enough.

**Figure 4:**
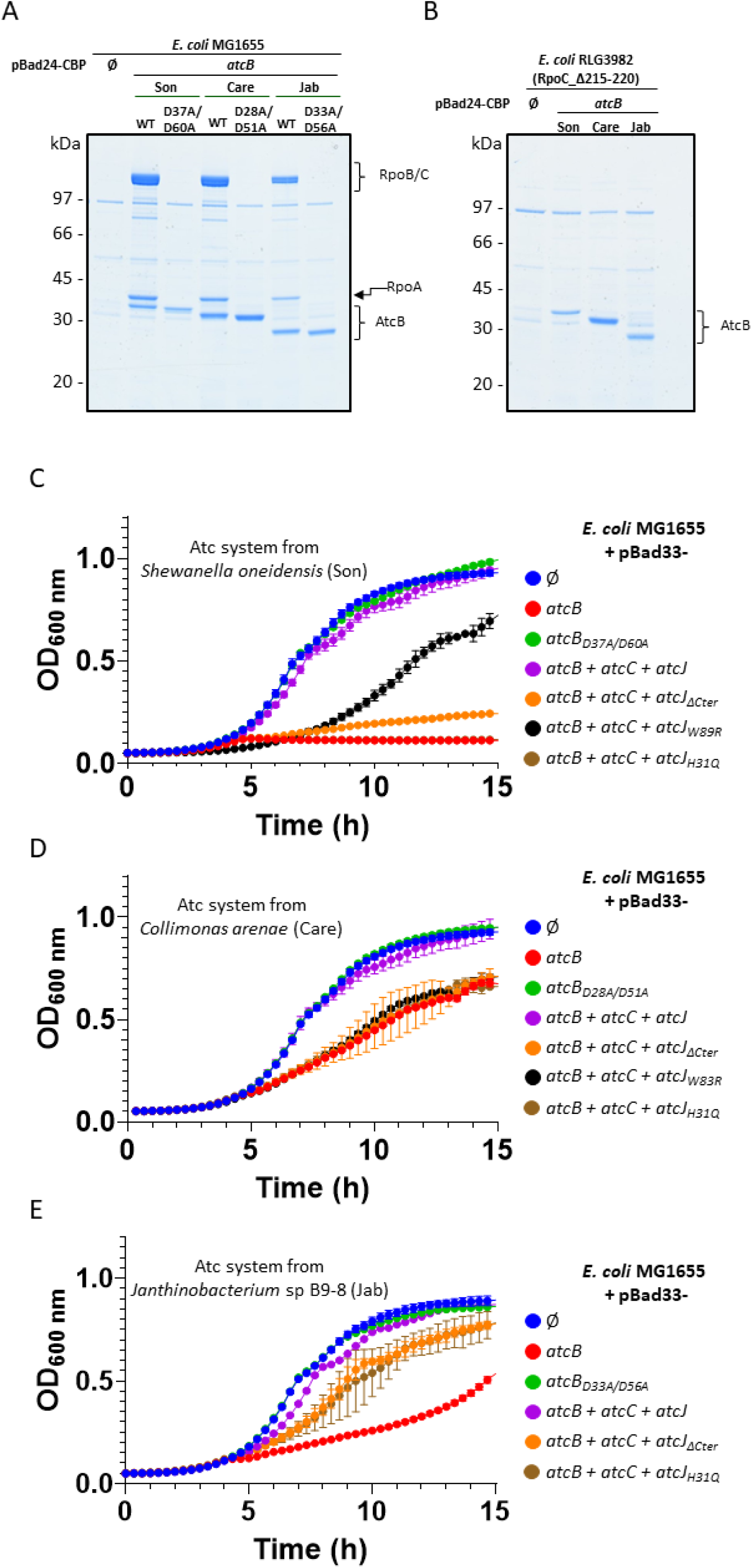
Molecular basis of AtcB–RNAP interaction and DnaK-mediated toxicity suppression. (A, B) In vivo co-purification of AtcB (WT or mutants) and RNAP in *E. coli* MG1655 (A) or RLG3982 (B) strains. Strains containing the pBad24 vector allow the overproduction of AtcB from *Shewanella oneidensis, Collimonas arenae* Cal35 and *Janthinobacterium* sp. B9-8 with a CBP-tag were grown at 37°C to the exponential growth phase before 0.2% arabinose was added for 2h. Control experiments with CBP produced alone were included (∅=empty vector). CBP-AtcB was purified on calmodulin beads. CBP-AtcB and the co-purified proteins were separated by SDS-PAGE, and stained with Coomassie blue. (C,D,E) Growth of *E. coli* MG1655 at 28°C producing from pBAD33 AtcB alone or in the presence of AtcC and AtcJ (WT or mutated) from *Shewanella oneidensis* (C)*, Collimonas arenae* Cal35 (D) and *Janthinobacterium* sp. B9-8 (E). Cultures were conducted in LB medium in the presence of chloramphenicol and 0.5% arabinose. After initial overnight growth at 28°C, strains were diluted to OD_600_ = 0.01, and were incubated at 28°C with shaking. Absorbance was measured over time. The data from two replicates are shown as mean ± SD. Note that in panel 4C, the growth curves of *E. coli* producing AtcB_Son_ either alone (red curve) or in the presence of AtcC and AtcJ_H31Q_ (brown curve) overlap.

We demonstrated before that AtcB from *S. oneidensis* interacts with AtcC using bacterial two-hybrid assays and co-immobilization experiments (Maillot *et al*. 2019; Boussouar *et al*. 2025). We aimed to confirm this interaction for the Care and Jab proteins as well. To test this, AtcB proteins were co-expressed in *E. coli* as CBP-tagged fusions alongside 6His-tagged AtcC. Cell extracts were then incubated with calmodulin resin to capture AtcB. After washings, AtcB and its associated proteins were eluted, separated by SDS-PAGE, and AtcC was detected by western blot using an anti-6His antibody. As for the Son system (**Figure 5A**), AtcB from Care and Jab co-eluted with their AtcC partner, supporting specific interaction (**Figure 5B**). Notably, AtcC proteins from Son and Care, when produced alone, bind weakly to calmodulin resin. However, the amount of detected AtcC increases substantially in the presence of CBP-AtcB, strongly supporting an interaction between the two proteins. Unfortunately, BACTH studies ran with the Care and Jab AtcB and AtcC proteins were unsuccessful since no β-galactosidase activity could be measured, contrary to the Son system. Instability of one or both fusion proteins, or non-productive AtcB-AtcC interaction preventing the spatial proximity of the T18 and T25 domains are possible explanations.

**Figure 5:**
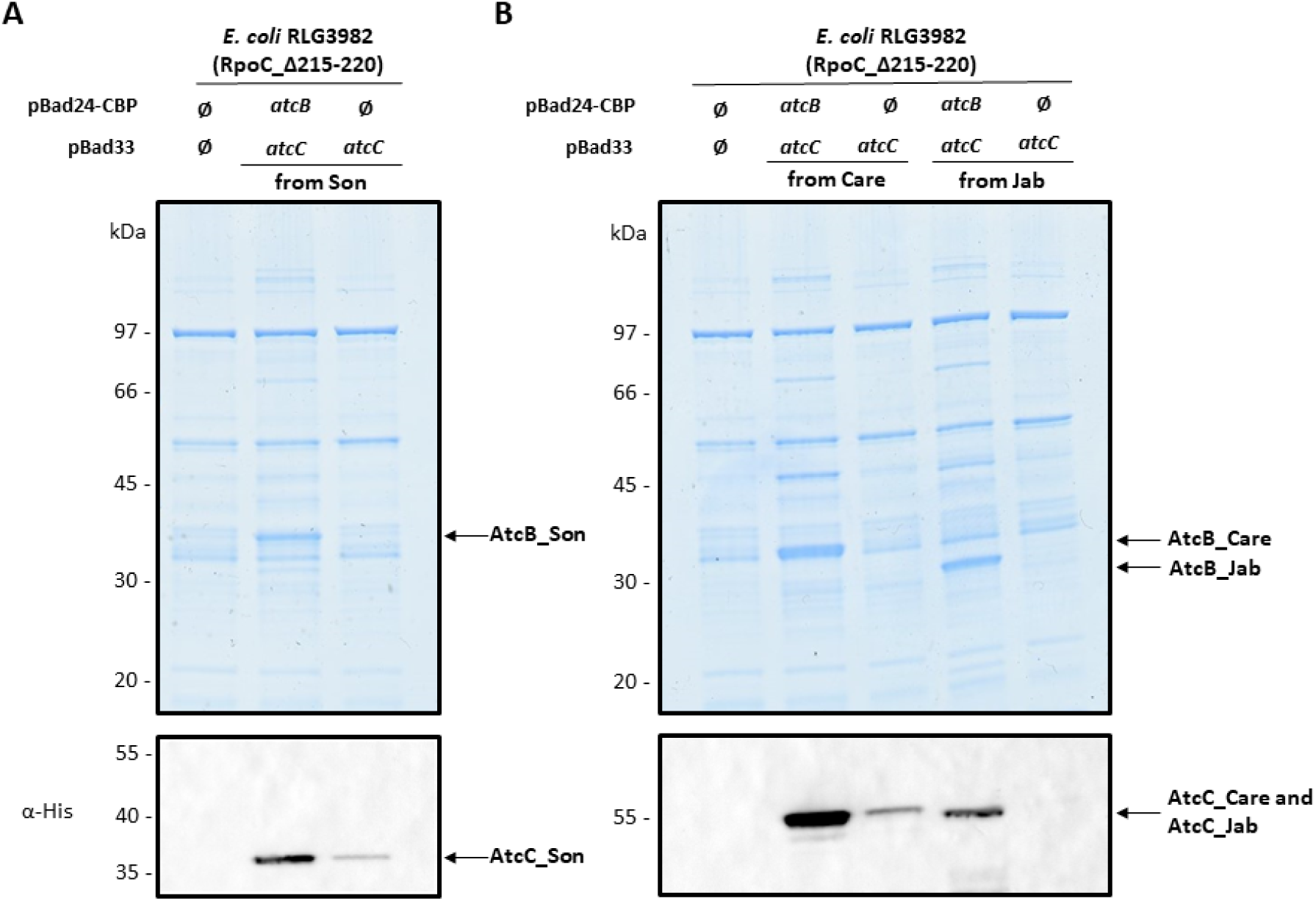
*Shewanella oneidensis*, *Collimonas arenae* Cal35 and *Janthinobacterium* sp. B9-8 AtcB-AtcC interactions assessed by pull-down assay. In vivo co-purification of AtcB and AtcC in *E. coli* RLG3982 (RpoC_Δ215-220). (A) *Shewanella oneidensis* (Son) proteins and (B) *Collimonas arenae* Cal35 and *Janthinobacterium* sp. B9-8 proteins. Cells containing the pBad24 vector either empty (Ø) or allowing the overproduction of AtcB with a CBP-tag, in the presence of pBad33 allowing the overproduction of AtcC-6His were grown at 37°C to the exponential growth phase before 0.2% arabinose was added for 2h. A control experiment with CBP produced alone was included. CBP-AtcB was purified on calmodulin beads. CBP-AtcB and the co-purified proteins were separated by SDS-PAGE, and stained with Coomassie blue. The presence of AtcC-6His as AtcB co-elutant was confirmed by Western blot using an anti-6His antibody.

Overall, these experiments support that the AtcJ-AtcC and AtcB-AtcC interactions are common markers of Atc systems, despite notable differences such as the lack of conservation of the AtcJ PX_7_W motif and the size of AtcC.

### Characterization of the Atc system from *Solidesulfovibrio magneticus* (Dma)

As shown above, the AtcB proteins from the Tsy, Veg, and Dma systems did not bind *E. coli* RNAP strongly enough to co-immobilize or cause toxicity (**Figure 3**). Yet, by focusing on the Dma system, we sought to determine whether the other interactions (AtcJ-AtcC and AtcB-AtcC), observed in the other systems, are also present in the Dma system. To address this, we employed bacterial two-hybrid (BACTH) assays and co-immobilization experiments. BACTH analysis confirmed that Dma AtcJ and AtcC interact, and that this interaction is specifically mediated by the C-terminal extension of AtcJ although it does not contain the PX_7_W (**Figure 6A**). Indeed, β-galactosidase activity was detected when using the C-terminal peptide alone, contrary to a truncated form of AtcJ lacking its C-terminal peptide. To test whether this interaction, like in the Son system, relies on hydrophobic contacts, we generated point mutants in which each aromatic hydrophobic residue of the C-terminal extension was individually substituted with serine. Surprisingly, none of these mutants showed reduced interaction. This suggests that the interaction may not depend on hydrophobic contacts, or that the binding is the result of a collective effect from the mutated residues. Further studies are needed to clarify this mechanism. The AlphaFold 3 structural model of the AtcJ–AtcC complex, shown in **Figure S1F**, supports the collective involvement of these hydrophobic residues despite its low confidence. The AtcB-AtcC interaction was also confirmed by both BACTH assay and co-immobilization experiments (**Figure 6B and 6C**).

**Figure 6:**
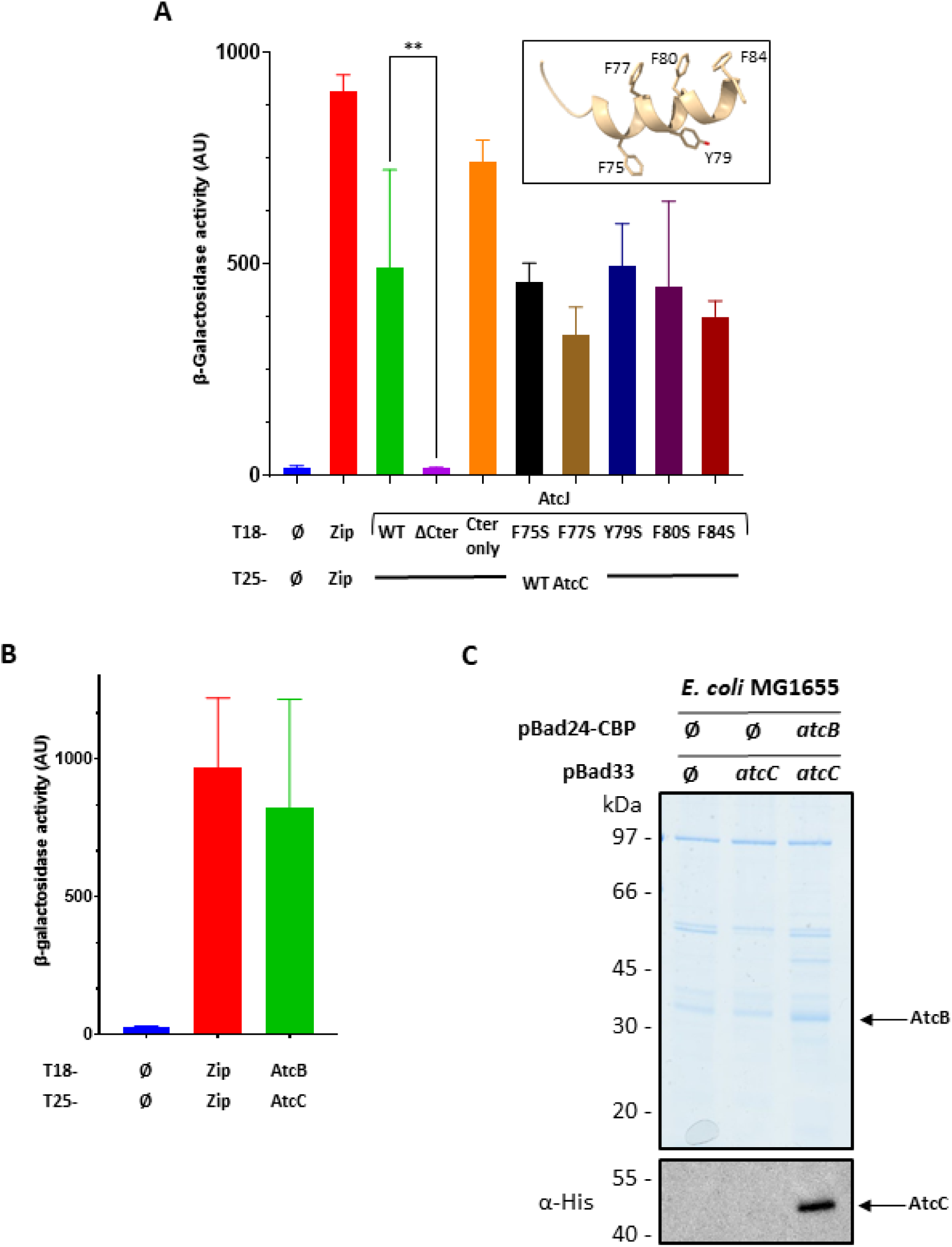
*Solidesulfovibrio magneticus* RS1 AtcJ-AtcC and AtcB-AtcC interactions assessed by bacterial two-hybrid assay and pull-down assay. (A) The *E. coli* BTH101 strain was co-transformed with the T18 and T25 plasmids coding for protein fusion between T18 and AtcJ (WT and mutants), A) or AtcB (B), and protein fusion between T25 and AtcC. The mutations in AtcJ are located in the C-terminal peptide, which is modeled as an alpha helix, specifically at the hydrophobic residues shown in the inset of panel A. A positive control (Zip/Zip, pUT18C-zip with pKT25-zip) and a negative control (Ø/Ø, empty vectors) were also included in the experiments. Cells were grown overnight at 28°C in the presence of IPTG. β-galactosidase activity was measured as explained in the Methods section. The data from three replicates are shown as mean ± SD and are analysed using a one-way ANOVA statistical test. P-values : ** <0.01, absence of labeling is for non-significant difference. (C) In vivo co-purification of AtcB and AtcC in *E. coli* MG1655. Strains containing the pBad24 vector either empty (Ø) or allowing the overproduction of AtcB with a CBP-tag, in the presence of pBad33 allowing the overproduction of AtcC-6His were grown at 37°C to the exponential growth phase before 0.2% arabinose was added for 2h. A control experiment with CBP produced alone was included. CBP-AtcB was purified on calmodulin beads. CBP-AtcB and the co-purified proteins were separated by SDS-PAGE, and stained with Coomassie blue. The presence of AtcC-6His as AtcB co-elutant was confirmed by Western blot using an anti-6His antibody.

### Phylogenetic analysis of the bacterial Atc systems

Following previous work (Maillot *et al*. 2019; Weber *et al*. 2023; Boussouar *et al*. 2025), we performed a more extensive analysis to determine which organisms possesses the *atc* operon using the available reference proteomes in Uniprot (UniProt Consortium 2025). The *atc* operon was identified in 294 genomes (**Supplementary File**), mainly in Beta- and Gammaproteobacteria and displayed a non-random phylogenetic distribution, with enrichment in specific clades associated with psychrophilic, mesophilic or aquatic lineages (**Figure 7**). No thermophilic bacteria were found possessing the *atc* operon. Interestingly, the *Janthinobacterium* sp. B9-8 strain used here is phylogenetically more closely related to an *Iodobacter* than to a *Janthinobacter*, as described previously (Xu *et al*. 2019). Both groups possess strains that are capable of producing violacein, which may explain their phylogenetic proximity. Interestingly, and unexpectedly, we also see here a group that appears monophyletic between *Telluria* and *Massilia*, which could suggest that these two genus should be grouped together, a possibility that has already been observed and suggested previously (Bowman 2023). Different phylogenetic trees were reconstructed for each component of the operon (see Materials and Methods) and compared to the species phylogenetic tree. The tanglegram analysis revealed strong overall congruence for AtcB, AtcC, and AtcJ which appear consistent with vertical inheritance, with relatively low entanglement values after optimization (AtcB, 0.008; AtcC, 0.024; AtcJ, 0.012) (see **Table S2**). In contrast, AtcA showed greater topological incongruence (entanglement = 0.256, **Figure S4 and Table S2**), which suggests increased lineage-specific diversification. Another correlation test was done with Mantel, by comparing phylogenetic distance matrices. AtcB, the RNAP-binding component, showed the strongest correlation with the species phylogeny (r = 0.867), followed by AtcC (r = 0.765) and AtcJ (r = 0.658), whereas AtcA was less strongly correlated (r = 0.442; all P < 10⁻⁴). The strong conservation of AtcB is consistent with its essential role in the interaction with the RNAP. In contrast, the AtcJ-AtcC module, which recruits DnaK to limit accumulation of inhibited polymerase, was somewhat more flexible while remaining broadly conserved. The strong phylogenetic conservation of AtcB, suggests that subtle lineage-specific differences at the AtcB-RNAP interface probably determine functional compatibility. By looking at pairwise evolutionary distances of operon proteins scaled with species phylogenetic distances, there are varying strengths across the components of the Atc operon. The RNAP interacting protein AtcB showed the strongest correlation (R² = 0.75), followed by AtcC (R² = 0.59) and the J-domain protein AtcJ (R² = 0.43), whereas AtcA showed substantially weaker correspondence (R² = 0.20) (**Table S2** and **Figure S5**). The adaptivity of chaperones like AtcJ and the co-interactor AtcC is consistent with a modulatory role mediated by DnaK. This is probably because chaperone systems are often adaptable to their environment and/or host. Interestingly, AtcB is under strong evolutionary constraint and closely follows host lineage divergence, which is probably determined by its interaction with the RNAP.

**Figure 7:**
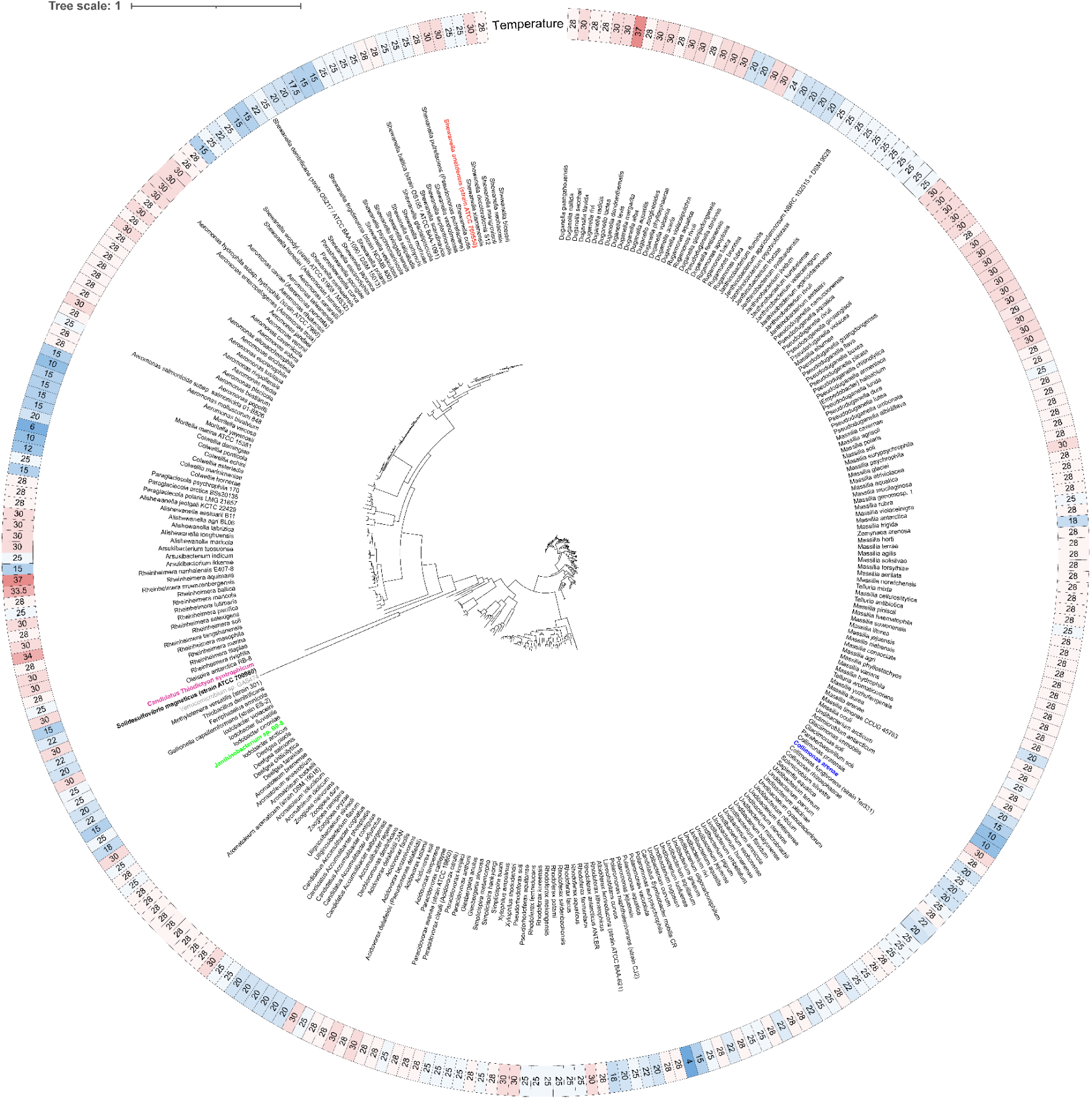
Circular species tree of 294 bacterial species possessing the *atc* operon with associated optimal growth temperature. The phylogenetic tree (constructed from 18 selected genes, see methods) depicts the evolutionary relationships among the studied species, including members of the genera *Shewanella*, *Aeromonas*, *Duganella*, *Massilia*, *Rhodoferax*, *Pseudoduganella*, *Janthinobacterium*, among others. The tree scale indicates the number of substitutions per site. The outer ring shows, for each species, the optimal growth temperature reported in the literature, expressed in °C. The color gradient, from blue (low temperatures, psychrophilic) to red (high temperatures, warm-mesophilic), illustrates the distribution of thermal preferences across the phylogeny. Selected species of interest are highlighted with specific colors: *Shewanella oneidensis* (red), *Candidatus Thiodictyon syntrophicum* (magenta), *Solidesulfovibrio magneticus* (bold), *Janthinobacterium sp. B9-8* (green), *Verrucomicrobium* sp. GAS474 (grey) and *Collimonas arenae* (blue).

## Discussion

By analyzing six Atc systems, we revealed a remarkable conservation of functional traits of the Atc system across environmental bacteria, despite significant sequence diversity in the key C-terminal motif of AtcJ (PX_7_W motif either degenerate or absent) or heterogeneity in AtcC size. The AtcJ-AtcC and AtcB-AtcC interactions were verified in all the tested cases. The conditional toxicity of AtcB provided critical functional insights into this system. When overexpressed in *E. coli*, AtcB proteins from *S. oneidensis*, *C. arenae*, and *Janthinobacterium* sp. B9-8 bind RNAP and probably inhibit transcription, resulting in growth defects. This toxicity is rescued by co-expression with AtcJ and AtcC, which recruit DnaK to prevent accumulation of the inhibitory AtcB-RNAP complex. The specificity of AtcB-RNAP interaction is governed by subtle structural determinants. Nevertheless, two conserved aspartate residues in AtcB (D37 and D60 in *S. oneidensis*) are essential for binding to a positively charged helix in the RpoC subunit of RNAP. The AtcB proteins from *Thyodiction syntrophicum*, *Verrucomicrobium* sp. GAS474, and *Solidesulfovibrio magneticus* fail to interact with *E. coli* RNAP because their structural interfaces may be incompatible. In the *atcB* gene tree, *Solidesulfovibrio magneticus* and *Verrucomicrobium* sp. GAS474 form a closely related pair, whereas *Thiodictyon syntrophicum* branches separately, nested among *Paraglaciecola*, *Colwellia*, and *Oleispira* sequences (**Figure S6**). This suggests that the incompatible AtcB structural interface may share a common origin in at least the *Solidesulfovibrio*/*Verrucomicrobium* pair, while its occurrence in *Thiodictyon syntrophicum* could reflect an independent structural divergence.

Phylogenetic analysis provides robust support for the functional conservation of the Atc system. The *atcB*, *atcC*, and *atcJ* genes exhibit strong congruence with species phylogeny, as evidenced by low entanglement values in tanglegram analysis (0.008, 0.024, and 0.012, respectively) and highly significant Mantel test correlations (r = 0.867, 0.765, and 0.658). These findings indicate vertical inheritance and co-evolution with the host’s transcriptional machinery. In striking contrast, *atcA* shows substantial topological incongruence (entanglement = 0.256, r = 0.442), suggesting a more flexible evolutionary trajectory and potentially distinct functional constraints.

Our results therefore highly suggest that the core interaction architecture (*ie* AtcJ-AtcC recruits the DnaK chaperone to modulate the inhibitory activity of AtcB on RNAP) remains functionally preserved even in non-canonical systems, validating a unified molecular mechanism. The enrichment of the Atc system in aquatic and psychrophilic lineages offers ecological context. Its complete absence in thermophiles suggests a specific adaptation to cold environments, where protein folding challenges and transcriptional regulation demands are heightened.

These findings support that the Atc system constitutes a new mechanism of transcriptional regulation that links chaperone protein activity to the control of gene expression, particularly under stress conditions. The vertical inheritance of this system suggests an ancient origin, while the observed sequence diversity reflects fine-tuned adaptation to different host contexts and environmental niches. Future work should focus on identifying the specific genes regulated by Atc, which could reveal new targets for the design of cold-adapted bacteria or the development of synthetic regulatory circuits.

## Acknowledgements

The authors thank the Bip01 team for helpful discussions. Martin Ardiley is acknowledged for performing preliminary experiments related to this study.

## Author contributions

Safa Boussouar (Conceptualization, Formal Analysis, Investigation, Visualization, Writing – review & editing), Olivier Genest (Supervision, Writing – review & editing, Funding acquisition), Mathieu E. Rebeaud (Formal Analysis, Investigation, Visualization, Writing – original draft, Writing – review & editing), Sébastien Dementin (Conceptualization, Formal Analysis, Investigation, Visualization, Supervision, Writing – original draft, Writing – review & editing, Project administration). The authors acknowledge that they used DeepL (https://www.deepl.com) and Mistral Vibe (https://chat.mistral.ai/chat) as translation tools while writing this article.

