## Supplementary Files for "Conserved Functional Traits of the Atc Protein System"

**Table S1**: Features of the Atc proteins of the selected Atc systems

| | AtcJ | AtcA | AtcB | AtcC | AtcJ-AtcC complex |
|---|---|---|---|---|---|
| ***Shewanella oneidensis* MR1 (Son)** | -94 aa<br>-C-ter motif :<br>**PX$_7$W (W89)**<br>-pTM=0.75 | -179 aa<br>-Potential IDP domain (D/E rich)<br>-pTM=0.6 | -252 aa<br>-Conserved Asp: D37 and D60<br>-pTM=0.72 | -298 aa<br>-pTM=0.82<br>-Two globular domains separated by long α-helix | -pTM=0.8<br>-ipTM=0.86<br>-Potential hydrophobic groove in N-terminal region of AtcC as interaction zone with C-ter peptide of AtcJ (W89 essential) |
| ***Collimonas arenae* Cal35 (Care)** | -92 aa<br>-Putative C-ter motif : **PX$_7$W (W83 involved in interaction with AtcC?)**<br>-pTM=0.7 | -182 aa<br>-Potential IDP domain (D/E rich)<br>-pTM=0.7 | -228 aa<br>-Conserved Asp: D28 and D51<br>-pTM=0.64 | -410 aa<br>-pTM=0.71<br>-Two globular domains separated by long α-helix | -pTM=0.6<br>-ipTM=0.59<br>-Potential hydrophobic groove in N-terminal region as interaction zone with C-ter peptide of AtcJ (W83 involved?) |
| ***Janthinobacterium* sp. B9-8 (Jab)** | -92 aa<br>-Putative C-ter motif : **PX$_7$A (I82, A83, Y86 involved in interaction with AtcC?)**<br>-pTM=0.73 | -178 aa<br>-Partially folded, C-terminal D/E rich domain<br>-pTM=0.7 | -254 aa<br>-Conserved Asp: D33 and D56<br>-pTM=0.78 | -405 aa<br>-pTM=0.64-Two globular domains separated by long α-helix | -pTM=0.58<br>-ipTM=0.55<br>-Potential hydrophobic groove in N-terminal region as interaction zone with C-ter peptide of AtcJ (I82, A83, Y86 involved?) |
| ***Thiodictyon syntrophicum* Cad16T (Tsy)** | -89 aa<br>-Putative C-ter motif: **AX$_7$W (W85 involved in interaction with AtcC?))**<br>-pTM=0.77 | -154 aa<br>-Potential IDP domain (D/E rich)<br>-pTM=0.77 | -237 aa<br>-Conserved Asp: D26 and D49<br>-pTM=0.73 | -394 aa<br>-pTM=0.18 | -pTM=0.35<br>-ipTM=0.51<br>-Potential hydrophobic groove in N-terminal region as interaction zone with C-ter peptide of AtcJ (W85 involved?) |
| ***Verrucomicrobium* sp. GAS474 (Veg)** | -97 aa<br>-C-ter : no conserved motif **(I81, Y84 involved in interaction with AtcC?))**<br>-pTM=0.64 | -136 aa<br>-No D/E rich C-ter region<br>-pTM=0.34 | -248 aa<br>-Conserved Asp: D39 and D62<br>-pTM=0.85 | -317 aa<br>-pTM=0.27 | -pTM=0.35<br>-ipTM=0.31 |
| ***Solidesulfovibrio magneticus* RS-1 (Dma)** | -85 aa<br>-C-ter : no conserved motif **(F75, F77, Y79, F80, F84 involved in interaction with AtcC?))**<br>-pTM=0.71 | -153 aa<br>-D/E rich Cter region<br>-pTM=0.19 | -257 aa<br>-Conserved Asp: D41 and D64<br>-pTM=0.79 | -408 aa<br>-pTM=0.25 | -pTM=0.19<br>-ipTM=0.29 |

**Table S2:** Statistics for the different tests of each protein (function, entanglement score, Mantel correlation (Spearman's r), and patristic distance comparison)

| Protein | Function | Mantel r | Patristic Distance $R^2$ | Entanglement |
|---|---|---|---|---|
| RpoC | RNAP core protein | 0.977 | 0.955 | 0.011 |
| DnaK | Hsp70 | 0.911 | 0.831 | 0.015 |
| AtcB | RNAP interaction | 0.867 | 0.751 | 0.008 |
| AtcC | adaptor/scaffold with DnaK ? | 0.765 | 0.585 | 0.024 |
| AtcJ | JDP | 0.658 | 0.432 | 0.012 |
| AtcA | divergent component / unknown activity | 0.442 | 0.195 | 0.256 |

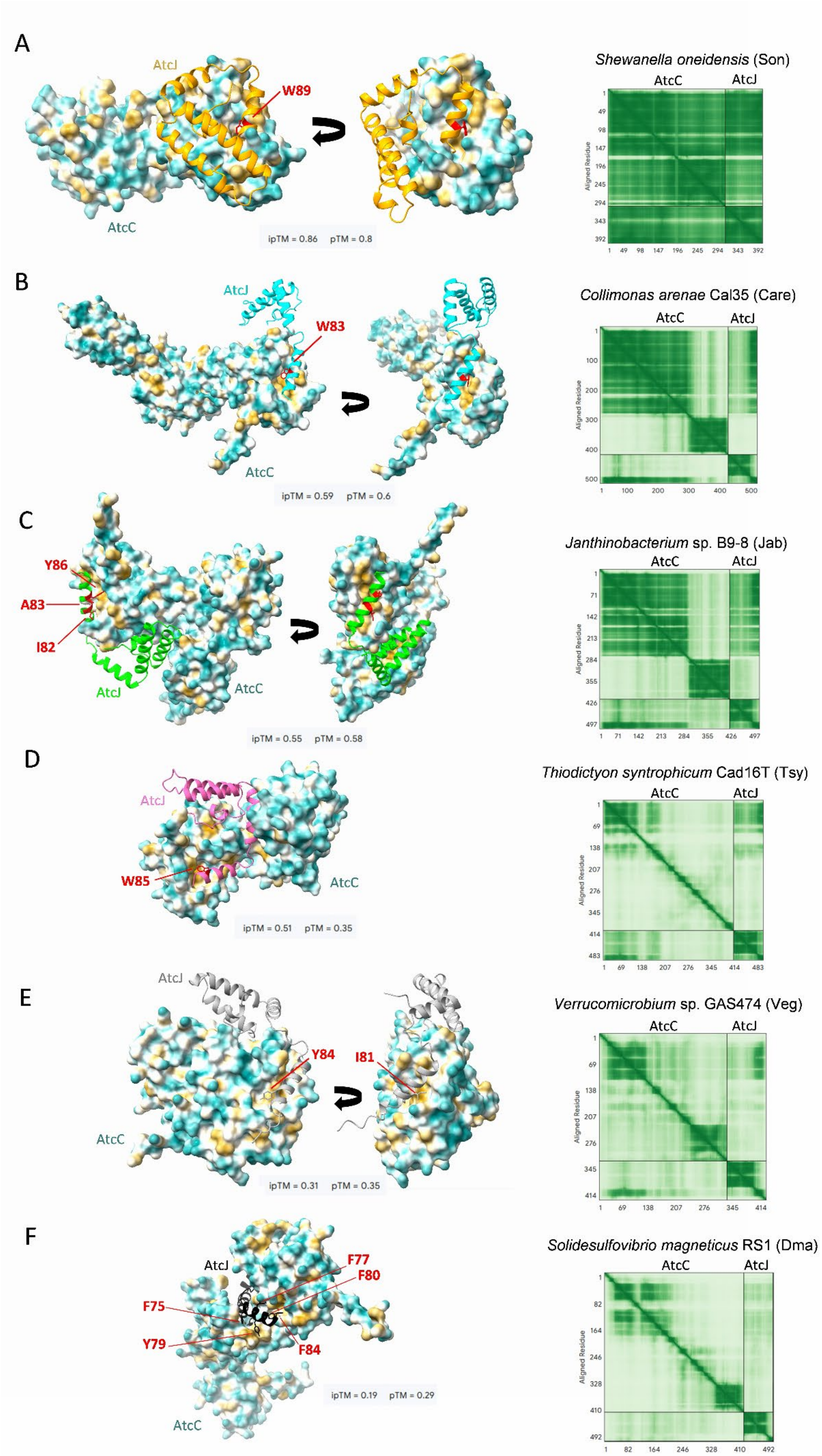
A
AtcJ
W89
AtcC
ipTM = 0.86 pTM = 0.8
Shewanella oneidensis (Son)
AtcC
AtcJ
Aligned Residue
B
AtcJ
W83
AtcC
ipTM = 0.59 pTM = 0.6
Collimonas arenae Cal35 (Care)
AtcC
AtcJ
C
Y86
A83
I82
AtcJ
AtcC
ipTM = 0.55 pTM = 0.58
Janthinobacterium sp. B9-8 (Jab)
AtcC
AtcJ
D
AtcJ
W85
AtcC
ipTM = 0.51 pTM = 0.35
Thiodictyon syntrophicum Cad16T (Tsy)
AtcC
AtcJ
E
AtcJ
Y84
I81
AtcC
ipTM = 0.31 pTM = 0.35
Verrucomicrobium sp. GAS474 (Veg)
AtcC
AtcJ
F
AtcJ
F77
F80
F75
Y79
F84
AtcC
ipTM = 0.19 pTM = 0.29
Solidesulfovibrio magneticus RS1 (Dma)
AtcC
AtcJ

**Figure S1: Structural models of the AtcJ-AtcC complexes from the selected organisms.** Structural predictions were obtained using the AlphaFold3 interface. The rank 1 structure models were selected and represented with ChimeraX software. Models of the complexes from (A) *Shewanella oneidensis* (Son), (B) *Collimonas arenae* Cal35 (Care), (C) *Janthinobacterium* sp. B9-8 (Jab), (D) *Thiodictyon syntrophicum* Cad16T (Tsy), (E) *Verrucomicrobium* sp. GAS474 (Veg) and (F) *Solidesulfovibrio magneticus* RS1 (Dma).

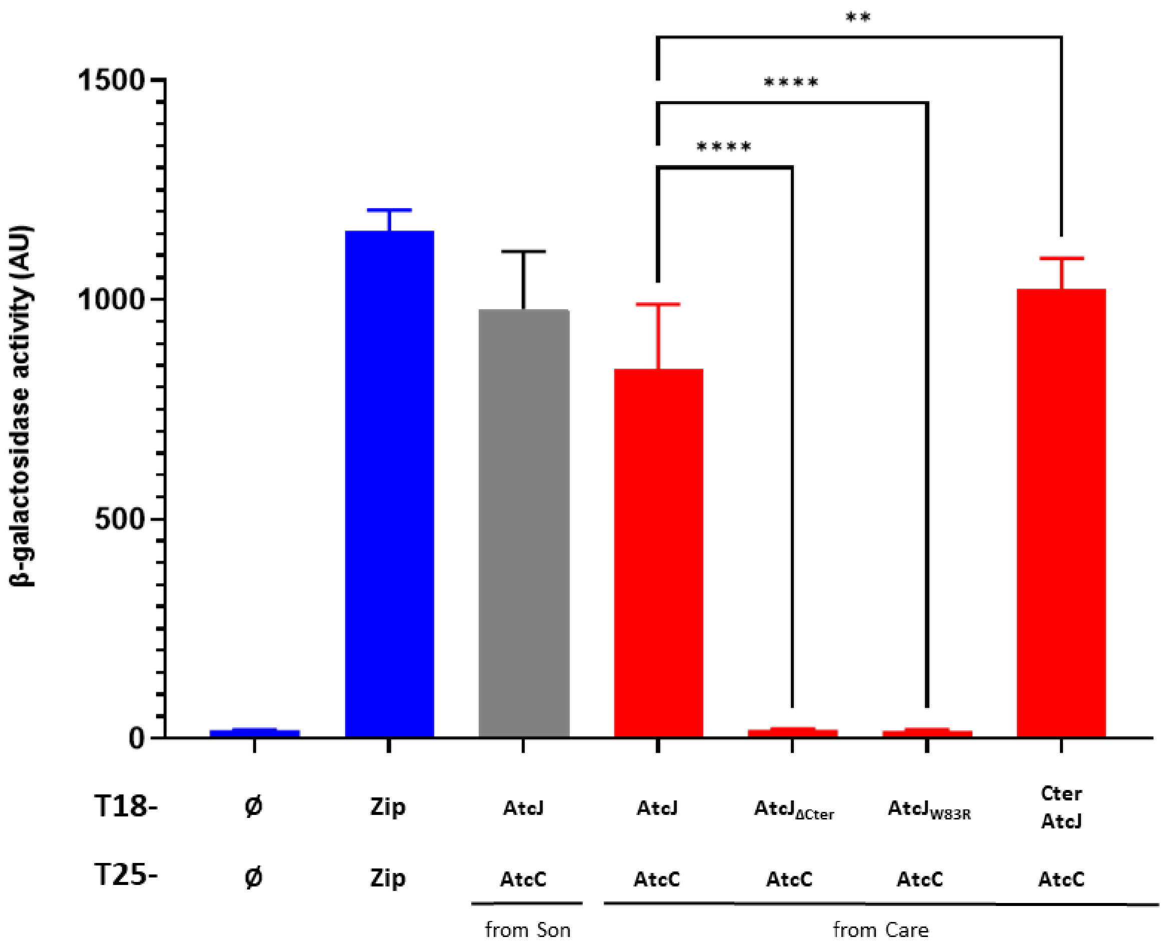


**Figure S2**: ***Shewanella oneidensis*** **(Son) and** ***Collimonas arenae*** **Cal35 (Care) AtcJ-AtcC interaction assessed by bacterial two-hybrid assay.** The *E. coli* BTH101 strain was co-transformed as indicated with the T18 and T25 plasmids coding for protein fusion between T18 and AtcJ (WT and mutants), and protein fusion between T25 and AtcC. A positive control (Zip/Zip, pUT18C-zip with pKT25-zip) and a negative control (Ø/Ø, empty vectors) were also included in the experiment. Cells were grown overnight at 28°C in the presence of IPTG. β-galactosidase activity was measured as explained in the Methods section. The data from three replicates are shown as mean ± SD and are analysed using a one-way ANOVA statistical test. P-values : **** <0.0001 and ** =0.0088, absence of labeling is for non-significant difference.

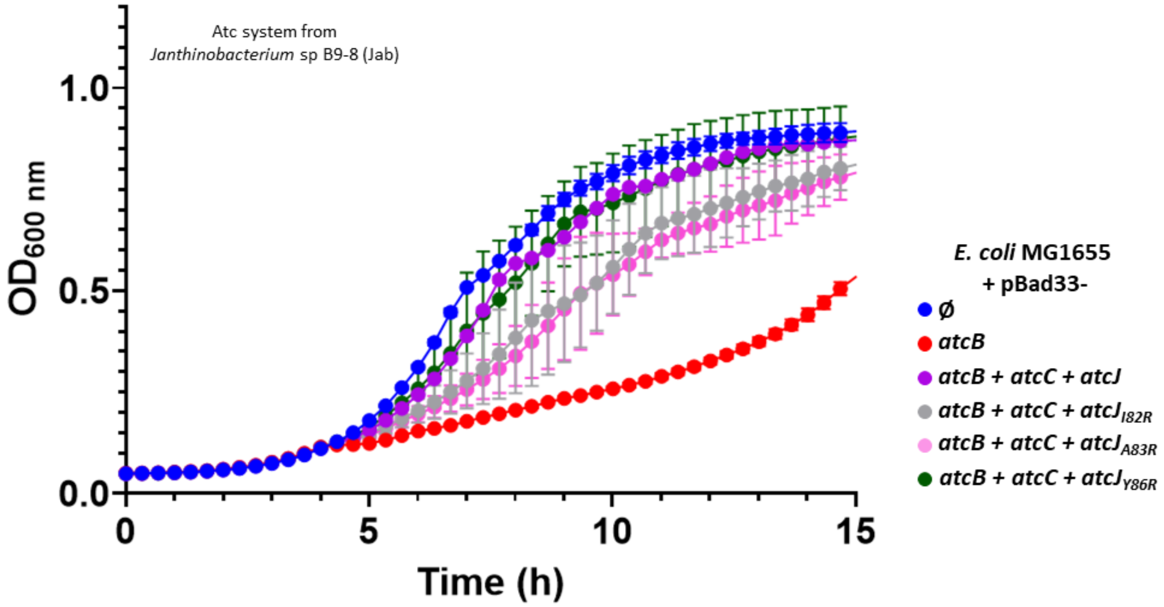


**Figure S3: Effect of mutations in *Janthinobacterium* sp. B9-8 AtcJ on the AtcJ-AtcC dependent alleviation of AtcB toxicity.** Growth of *E. coli* MG1655 at 28°C producing from pBad33 *Janthinobacterium* sp. B9-8 AtcB alone or in the presence of AtcC and AtcJ (WT or mutated) . Cultures were conducted in LB medium in the presence of chloramphenicol and 0.5% arabinose. After initial overnight growth at 28°C, strains were diluted to OD600nm = 0.01, and were incubated at 28°C with shaking. Absorbance was measured over time. The data from two replicates are shown as mean ± SD.

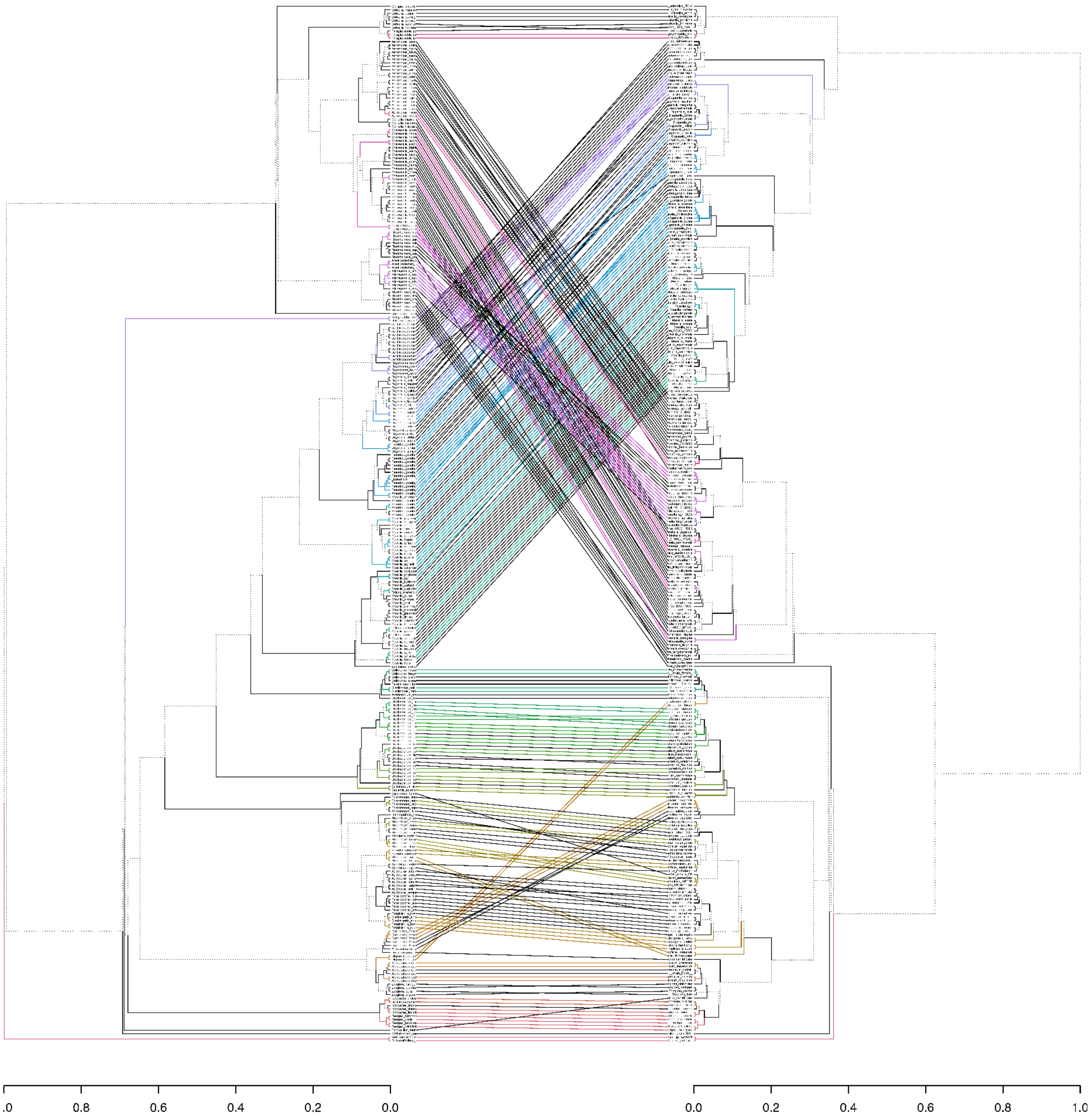


**Figure S4: Tanglegram comparing the species phylogeny with AtcA tree.** The left tree is the species phylogeny, and the right tree is inferred from the *atcA* gene sequence, plotted using the R package dendextend and ape. The tip order in both trees was optimized using the step2side untangling algorithm. The branches shared as common subtrees between the two trees are colored consistently, while branches not part of a common subtree are shown as dotted black lines. Extensive crossing of connecting lines reflects substantial topological incongruence between the *atcA* gene tree and the species tree. The corresponding entanglement score is reported in Supplementary Table 2.

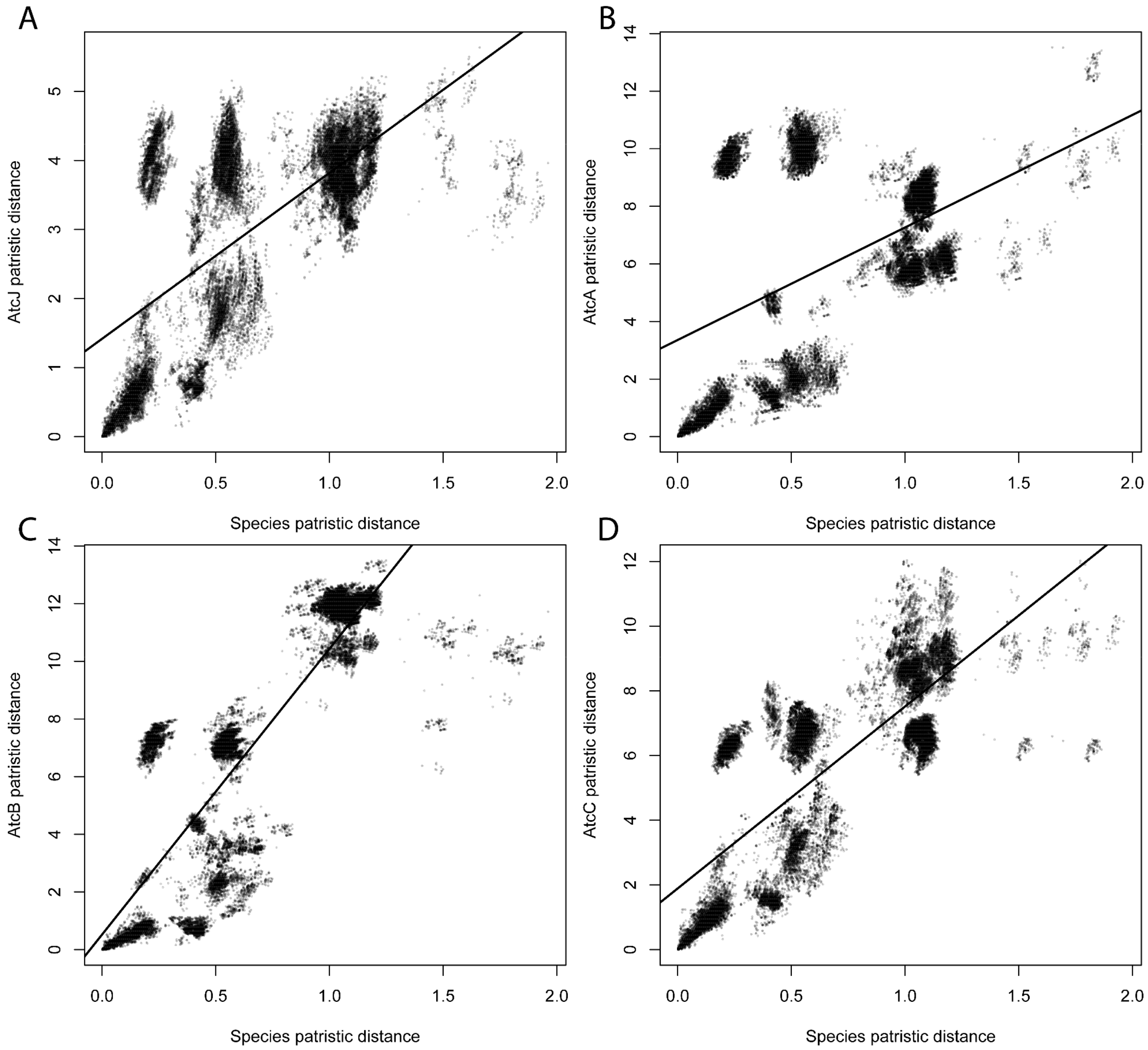


**Figure S5: Patristic distance between species tree and *atc* gene family (*atcA*, *atcB*, *atcC*, *atcJ*).** Each panel plots patristic distances derived from the phylogenetic tree of each gene against the species tree. (A) *atcJ* vs. species patristic distance. (B) *atcA* vs. species patristic distance. (C) *atcB* vs. species patristic distance. (D) *atcC* vs. species patristic distance. In each panel, the solid black line represents the linear regression fit between gene and species distances. Deviations from the fitted line, and points falling above or below the main trend, indicate differential rates of sequence divergence of the *atc* genes relative to overall species divergence. The corresponding patristic distance comparison is reported in Supplementary Table 2.

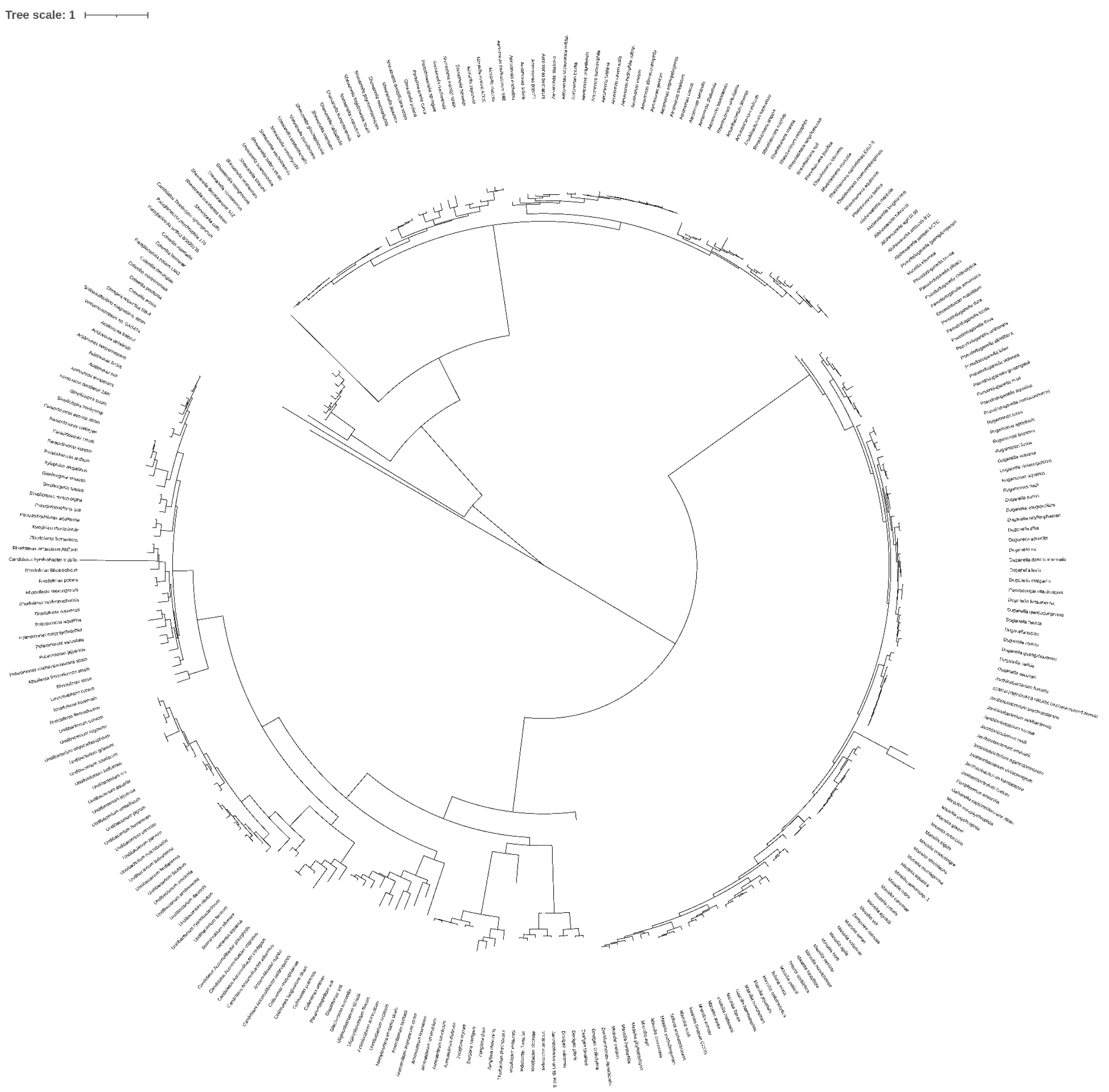


**Figure S6: Circular *atcB* gene tree of the 294 bacterial species.** The tree was inferred from AtcB protein sequences using IQ-TREE under the LG+G4+I+F substitution model. Branch lengths represent the expected number of amino acid substitutions per site. The tree includes representatives from genera such as *Acidovorax*, *Rhodoferax*, *Undibacterium*, *Aromatoleum*, *Zoogloea*, *Iodobacter*, *Janthinobacterium*, *Duganella*, *Pseudoduganella*, *Massilia*, *Rheinheimera*, *Alishewanella*, *Aeromonas*, *Shewanella*, *Colwellia*, and *Paraglaciecola*, among others.

**Supplementary Material**

Table S1: Strains used in this study

| Strains of *E. coli* | Genotype | Reference |
|---|---|---|
| C600 | F- *tonA21 thi-1 thr-1 leuB6 lacY1 glnV44 rfbC1 fhuA1 λ-* | (Appleyard 1954) |
| MG1655 | F- lambda- *ilvG- rfb*-50 *rph*-1 | (Blattner *et al.* 1997) |
| RLG3982 (RpoC_Δ215-220) | MG1655 *lac*Δ145 *thi39*::Tn10 *rpoC* (β' Δ215-220) λRLG957<br><br>λRLG957 contains *rrnB* P1 (-61 to +50)-*lacZ* | (Bartlett *et al.* 1998) |
| BTH101 | F- *cya-99 araD139 galE15 galK16 rpsL1* ($Str^R$) *hsdR2 mcrA1 mcrB1* | (Battesti and Bouveret 2012) |

Table S2: Plasmids used in this study

| Plasmid | Description | Reference |
|---|---|---|
| pBad33 | Vector for arabinose inducible expression containing PBad promoter with the p15A origin of replication, $Cm^R$ | (Guzman *et al.* 1995) |
| pBad33-*atcJ_Son* | Sequence coding for AtcJ from *Shewanellla oneidensis* cloned into pBad33 | (Maillot *et al.* 2019) |
| pBad33-*atcC_Son* | Sequence coding for AtcC from *Shewanellla oneidensis* 6His-tagged in C-ter cloned into pBad33 | (Maillot *et al.* 2019) |
| pBad33-*atcC* + *atcJ_Son* | Sequences coding for AtcC 6His-tagged and AtcJ cloned from *Shewanellla oneidensis* into pBad33 | (Boussouar *et al.* 2025) |
| pBad33-*atcB_Son* | Sequence coding for AtcB from *Shewanellla oneidensis* 6His-tagged in C-ter cloned into pBad33 | (Maillot *et al.* 2019) |
| pBad33-*atcB_Son*$_{D37A/D60A}$ | pBad33-*atcB* from *Shewanellla oneidensis* containing mutations to produce D37A/D60A AtcB variant | (Boussouar *et al.* 2025) |
| pBad33-*atcB* + *atcC* + *atcJ_Son* | Sequences coding for AtcB 6His-tagged in C-ter, AtcC 6His-tagged in C-ter and AtcJ from *Shewanellla oneidensis* cloned into pBad33 | (Boussouar *et al.* 2025) |
| pBad33-*atcB* + *atcC* + *atcJ_Son*$_{\Delta Cter}$ | pBad33-*atcB* + *atcC* + *atcJ* from *Shewanellla oneidensis* with mutation to produce ΔCter AtcJ variant | (Boussouar *et al.* 2025) |
| pBad33-*atcB* + *atcC* + *atcJ_Son*$_{H31Q}$ | pBad33-*atcB* + *atcC* + *atcJ* from *Shewanellla oneidensis* with mutation to produce H31Q AtcJ variant | (Boussouar *et al.* 2025) |
| pBad33-*atcB* + *atcC* + *atcJ_Son*$_{W89R}$ | pBad33-*atcB* + *atcC* + *atcJ* from *Shewanellla oneidensis* with mutation to produce W89R AtcJ variant | This work |
| pBad33-*atcJ_Care* | Sequence coding for AtcJ from *Collimonas arenae* Cal35 cloned into pBad33 | This work |
| pBad33-*atcC_Care* | Sequence coding for AtcC from *Collimonas arenae* Cal35 6His-tagged in C-ter cloned into pBad33 | This work |
| pBad33-*atcC* + *atcJ_Care* | Sequences coding for AtcC 6His-tagged and AtcJ cloned from *Collimonas arenae* Cal35 into pBad33 | This work |
| pBad33-*atcB_Care* | Sequence coding for AtcB from *Collimonas arenae* Cal35 6His-tagged in C-ter cloned into pBad33 | This work |

| | | |
|---|---|---|
| pBad33-*atcB_Care*$_{D28A/D51A}$ | pBad33-*atcB* from *Collimonas arenae* Cal35 containing mutations to produce D28A/D51A AtcB variant | This work |
| pBad33-*atcB* + *atcC* + *atcJ_Care* | Sequences coding for AtcB 6His-tagged in C-ter, AtcC 6His-tagged in C-ter and AtcJ from *Collimonas arenae* Cal35 cloned into pBad33 | This work |
| pBad33-*atcB* + *atcC* + *atcJ_Care*$_{\Delta Cter}$ | pBad33-*atcB* + *atcC* + *atcJ* from *Collimonas arenae* Cal35 with mutation to produce ΔCter AtcJ variant | This work |
| pBad33-*atcB* + *atcC* + *atcJ_Care*$_{H31Q}$ | pBad33-*atcB* + *atcC* + *atcJ* from *Collimonas arenae* Cal35 with mutation to produce H31Q AtcJ variant | This work |
| pBad33-*atcB* + *atcC* + *atcJ_Care*$_{W83R}$ | pBad33-*atcB* + *atcC* + *atcJ* from *Collimonas arenae* Cal35 with mutation to produce W83R AtcJ variant | This work |
| pBad33-*atcJ_Jab* | Sequence coding for AtcJ from *Janthinobacterium* sp. B9-8 cloned into pBad33 | This work |
| pBad33-*atcC_ Jab* | Sequence coding for AtcC from *Janthinobacterium* sp. B9-8 6His-tagged in C-ter cloned into pBad33 | This work |
| pBad33-*atcC* + *atcJ_Jab* | Sequences coding for AtcC 6His-tagged and AtcJ cloned from *Janthinobacterium* sp. B9-8 into pBad33 | This work |
| pBad33-*atcB_ Jab* | Sequence coding for AtcB from *Janthinobacterium* sp. B9-8 6His-tagged in C-ter cloned into pBad33 | This work |
| pBad33-*atcB_ Jab* $_{D33A/D56A}$ | pBad33-*atcB* from *Janthinobacterium* sp. B9-8 containing mutations to produce D33A/D56A AtcB variant | This work |
| pBad33-*atcB* + *atcC* + *atcJ_ Jab* | Sequences coding for AtcB 6His-tagged in C-ter, AtcC 6His-tagged in C-ter and AtcJ from *Janthinobacterium* sp. B9-8 cloned into pBad33 | This work |
| pBad33-*atcB* + *atcC* + *atcJ_ Jab*$_{\Delta Cter}$ | pBad33-*atcB* + *atcC* + *atcJ* from *Janthinobacterium* sp. B9-8 with mutation to produce ΔCter AtcJ variant | This work |
| pBad33-*atcB* + *atcC* + *atcJ_ Jab* $_{H31Q}$ | pBad33-*atcB* + *atcC* + *atcJ* from *Janthinobacterium* sp. B9-8 with mutation to produce H31Q AtcJ variant | This work |
| pBad33-*atcB* + *atcC* + *atcJ_ Jab* $_{I82R}$ | pBad33-*atcB* + *atcC* + *atcJ* from *Collimonas arenae* with mutation to produce I82R AtcJ variant | This work |

| | | |
|---|---|---|
| pBad33-*atcB* + *atcC* + *atcJ_ Jab* $_{Y86R}$ | pBad33-*atcB* + *atcC* + *atcJ* from *Collimonas arenae* with mutation to produce Y86R AtcJ variant | This work |
| pBad33-*atcB_Veg* | Sequence coding for AtcB from *Verrucomicrobium* sp. GAS474 6His-tagged in C-ter cloned into pBad33 | This work |
| pBad33-*atcB_Tsy* | Sequence coding for AtcB from *Thiodictyon syntrophicum* Cad16T 6His-tagged in C-ter cloned into pBad33 | This work |
| pBad33-*atcB_Dma* | Sequence coding for AtcB from *Solidesulfovibrio magneticus* RS1 6His-tagged in C-ter cloned into pBad33 | This work |
| pBad33-*atcC_ Jab* | Sequence coding for AtcC from *Janthinobacterium* sp. B9-8 6His-tagged in C-ter cloned into pBad33 | This work |
| pBad24-CBP-linker | Sequence coding for the calmodulin-binding protein (CBP) cloned into pBad24, $Amp^R$ | (Battesti and Bouveret 2008) |
| pBad24-CBP-*atcB_Son* | Sequence coding for AtcB from *Shewanellla oneidensis* cloned into pBad24-CBP-linker | (Maillot *et al.* 2021) |
| pBad24-CBP-*atcB_Son* $_{D37A/D60A}$ | pBad24-CBP-*atcB* from *Shewanellla oneidensis* with mutation to produce CBP-AtcB $_{D37A/D60A}$ | (Boussouar *et al.* 2025) |
| pBad24-CBP-*atcB_Care* | Sequence coding for AtcB from *Collimonas arenae* Cal35 cloned into pBad24-CBP-linker | This work |
| pBad24-CBP-*atcB_Care* $_{D28A/D51A}$ | pBad24-CBP-*atcB* from *Collimonas arenae* Cal35 with mutation to produce CBP-AtcB $_{D28A/D51A}$ | This work |
| pBad24-CBP-*atcB_Jab* | Sequence coding for AtcB from *Janthinobacterium* sp. B9-8 cloned into pBad24-CBP-linker | This work |
| pBad24-CBP-*atcB_Jab* $_{D33A/D56A}$ | pBad24-CBP-*atcB* from *Janthinobacterium* sp. B9-8 with mutation to produce CBP-AtcB $_{D33A/D56A}$ | This work |
| pBad24-CBP-*atcB_Veg* | Sequence coding for AtcB from *Verrucomicrobium* sp. GAS474 cloned into pBad24-CBP-linker | This work |
| pBad24-CBP-*atcB_Tsy* | Sequence coding for AtcB from *Thiodictyon syntrophicum* Cad16T cloned into pBad24-CBP-linker | This work |
| pBad24-CBP-*atcB_Dma* | Sequence coding for AtcB from *Solidesulfovibrio magneticus* RS1 cloned into pBad24-CBP-linker | This work |

| | | |
|---|---|---|
| pT18-*atcJ_Son* | *atcJ* sequence from *Shewanellla oneidensis* cloned in-frame at the 3′ end of the sequence coding for the T18 domain into pUT18-C-linker vector | (Maillot *et al.* 2019) |
| pT18-*atcB_Son* | *atcB* sequence from *Shewanellla oneidensis* cloned in-frame at the 3′ end of the sequence coding for the T18 domain into pUT18-C-linker vector | (Maillot *et al.* 2019) |
| pT25-*atcC_Son* | *atcC* sequence from *Shewanellla oneidensis* cloned in-frame at the 3′ end of the sequence coding for the T25 domain into pKT25-linker vector | (Maillot *et al.* 2019) |
| pT18-*atcJ_Care* | *atcJ* sequence from *Collimonas arenae* Cal35 cloned in-frame at the 3′ end of the sequence coding for the T18 domain into pUT18-C-linker vector | This work |
| pT18-*atcB_Son* | *atcB* sequence from *Collimonas arenae* cal35 cloned in-frame at the 3′ end of the sequence coding for the T18 domain into pUT18-C-linker vector | This work |
| pT25-*atcC_Care* | *atcC* sequence from *Collimonas arenae* Cal35 cloned in-frame at the 3′ end of the sequence coding for the T25 domain into pKT25-linker vector | This work |
| pT18-*atcJ_Jab* | *atcJ* sequence from *Janthinobacterium* sp. B9-8 cloned in-frame at the 3′ end of the sequence coding for the T18 domain into pUT18-C-linker vector | This work |
| pT18-Cter*atcJ_Jab* | Coding sequence of the Cter peptide of AtcJ from *Janthinobacterium* sp. B9-8 cloned in-frame at the 3′ end of the sequence coding for the T18 domain into pUT18-C-linker vector | This work |
| pT18-*atcB_Jab* | *atcB* sequence from *Janthinobacterium* sp. B9-8 cloned in-frame at the 3′ end of the sequence coding for the T18 domain into pUT18-C-linker vector | This work |

| | | |
|---|---|---|
| pT25-*atcC_jab* | *atcC* sequence from *Janthinobacterium* sp. B9-8 cloned in-frame at the 3′ end of the sequence coding for the T25 domain into pKT25-linker vector | This work |
| pT18-*atcJ_Dma* | *atcJ* sequence from *Solidesulfovibrio magneticus* RS1 cloned in-frame at the 3′ end of the sequence coding for the T18 domain into pUT18-C-linker vector | This work |
| pT18-*atcJ_Dma*$_{F75S}$ | pT18-*atcJ_Dma* with mutation to produce F75S AtcJ variant | This work |
| pT18-*atcJ_Dma*$_{F77S}$ | pT18-*atcJ_Dma* with mutation to produce F77S AtcJ variant | This work |
| pT18-*atcJ_Dma*$_{Y79S}$ | pT18-*atcJ_Dma* with mutation to produce Y79S AtcJ variant | This work |
| pT18-*atcJ_Dma*$_{F80S}$ | pT18-*atcJ_Dma* with mutation to produce F80S AtcJ variant | This work |
| pT18-*atcJ_Dma*$_{F84S}$ | pT18-*atcJ_Dma* with mutation to produce F84S AtcJ variant | This work |
| pT18-*atcJΔCter_Dma* | *atcJ* sequence from *Solidesulfovibrio magneticus* RS1 deleted from sequence coding for Cter peptide cloned in-frame at the 3′ end of the sequence coding for the T18 domain into pUT18-C-linker vector | This work |
| pT18-*CteratcJ_Dma* | Coding sequence of the Cter peptide of AtcJ from *Solidesulfovibrio magneticus* RS1 cloned in-frame at the 3′ end of the sequence coding for the T18 domain into pUT18-C-linker vector | This work |
| pT18-*atcB_Dma* | *atcB* sequence from *Solidesulfovibrio magneticus* RS1 cloned in-frame at the 3′ end of the sequence coding for the T18 domain into pUT18-C-linker vector | This work |
| pT25-*atcC_Dma* | *atcC* sequence from *Solidesulfovibrio magneticus* RS1 cloned in-frame at the 3′ end of the sequence coding for the T25 domain into pKT25-linker vector | This work |

Table S3: Primers used in this study

| Primer sequence (5' →3') | Usage |
|---|---|
| GGCTAGCGAATTCGAGCTCGAAGGAGATATACCATGAAGGATTATTACGCCGTT (f)<br>CGACTCTAGAGGATCCCCGGTTATTGCAAAACCTTGTTCATA (r) | To construct pBad33-*atcJ_Care*<br>Template: chromosome<br>The PCR product was cloned into PstI-linearized pBad33 using NEBuilder® HiFi DNA Assembly Cloning Kit. |
| GGCTAGCGAATTCGAGCTCGAAGGAGATATACCATGAAAGCAGCAAACCTGATCCGC (f)<br>GACTCTAGAGGATCCCCGGTTAGTGGTGATGATGGTGATGCAGGGCGGATTCCTCGC (r) | To construct pBad33-*atcB_Care*<br>Template: chromosome<br>The PCR product was cloned into PstI-linearized pBad33 using NEBuilder® HiFi DNA Assembly Cloning Kit. |
| GGCTAGCGAATTCGAGCTCGAAGGAGATATACCATGTCCGACAAGAACCGGCAT (f)<br>GACTCTAGAGGATCCCCGGTTAGTGGTGATGATGGTGATGTTTCTTTTCCTTGCCGGC (r) | To construct pBad33-*atcC_Care*<br>Template: chromosome<br>The PCR product was cloned into PstI-linearized pBad33 using NEBuilder® HiFi DNA Assembly Cloning Kit. |
| CGGGGATCCTCTAGAGTCGACCTCTACTGTTTCTCCATACCC (f)<br>AAACAGCCAAGCTTGCATGCCTTATTGCAAAACCTTGTTCA (r) | To construct pBad33-*atcC+ atcJ_Care*<br>Template: pBad33-*atcJ_Care*<br>The PCR product was cloned into PstI-linearized pBad33-*atcC_Care* using NEBuilder® HiFi DNA Assembly Cloning Kit. |
| CACCATTAACCGGGGATCCTTACCTGACGCTTTTTATCGC (f)<br>TGCATGCCTGCAGGTCGACTTTATTGCAAAACCTTGTTCA (r) | To construct pBad33-*atcB* + *atcC+ atcJ_Care*<br>Template: pBad33-*atcC+atcJ_Care*<br>The PCR product was cloned into XbaI-linearized pBad33-*atcB_Care* using NEBuilder® HiFi DNA Assembly Cloning Kit. |
| GGCTAGCGAATTCGAGCTCG (f)<br>GACTCTAGAGGATCCCCGG (r) | To construct pBad33-*atcJ_Jab*, pBad33-*atcB_Jab*, pBad33-*atcC_Jab*, pBad33-*atcB_veg*, pBad33-*atcB_Tsy*<br>Template: synthetic genes<br>The PCR product was cloned into PstI-linearized pBad33 using NEBuilder® HiFi DNA Assembly Cloning Kit. |

| | |
|---|---|
| CACCACTAACCGGGGATCCTCTCTACTGTTTCTCCATACCC (f)<br>TGCATGCCTGCAGGTCGACTTTACTGGATGATTCCTTTAATGT (r) | To construct pBad33-*atcC*+ *atcJ_jab*<br>Template: pBad33-*atcJ_Jab*<br>The PCR product was cloned into PstI-linearized pBad33-*atcC_Jab* using NEBuilder® HiFi DNA Assembly Cloning Kit. |
| CACCATTAACCGGGGATCCTTACCTGACGCTTTTTATCGC (f)<br>TGCATGCCTGCAGGTCGACTTTACTGGATGATTCCTTTAA (r) | To construct pBad33-*atcB* + *atcC*+ *atcJ_Jab*<br>Template: pBad33-*atcC*+ *atcJ_Jab*<br>The PCR product was cloned into XbaI-linearized pBad33-*atcB* *_Jab* using NEBuilder® HiFi DNA Assembly Cloning Kit. |
| GGCTAGCGAATTCGAGCTCGAAGGAGATATACCATGAATTATTACGATATTTTG (f)<br>CGACTCTAGAGGATCCCCGGTTATTTAAACAGAGTCAAGAA (r) | To construct pBad33-*atcJ_Dma*<br>Template: chromosome<br>The PCR product was cloned into PstI-linearized pBad33 using NEBuilder® HiFi DNA Assembly Cloning Kit. |
| GGCTAGCGAATTCGAGCTCGAAGGAGATATACCATGAAGTCTATTGCTTTGCCGAAG (f)<br>GACTCTAGAGGATCCCCGGTTAGTGGTGATGATGGTGATGGAGGTCTTCAAAAAGGT (r) | To construct pBad33-*atcB_Dma*<br>Template: chromosome<br>The PCR product was cloned into PstI-linearized pBad33 using NEBuilder® HiFi DNA Assembly Cloning Kit. |
| GGCTAGCGAATTCGAGCTCGAAGGAGATATACCATGACCTTTTTGAAGACCTCTACCC (f)<br>CGACTCTAGAGGATCCCCGGTTAGTGGTGATGATGGTGATGCCTACAATTCTTATGAA (r) | To construct pBad33-*atcC_Dma*<br>Template: chromosome<br>The PCR product was cloned into PstI-linearized pBad33 using NEBuilder® HiFi DNA Assembly Cloning Kit. |
| CGGGGATCCTCTAGAGTCGACCTCTACTGTTTCTCCATACCC (f)<br>AAACAGCCAAGCTTGCATGCCTTATTTAAACAGAGTCAAGA (r) | To construct pBad33-*atcC*+ *atcJ_Dma*<br>Template: pBad33-*atcJ_Dma*<br>The PCR product was cloned into PstI-linearized pBad33-*atcC_Dma* using NEBuilder® HiFi DNA Assembly Cloning Kit. |

| | |
|---|---|
| GGGCACTTGTCGAGTCGACTATGAAAGCAGCAAACCTGAT (f)<br>CCTGCAGGTCGAACTCGAGTTTACAGGGCGGATTCCTCGCCAT (r) | To construct pBad24-CBP-*atcB_Care*<br>Template: chromosome<br>The PCR product was cloned into XbaI-linearized pBad24-CBPlinker using NEBuilder® HiFi DNA Assembly Cloning Kit. |
| GGGCACTTGTCGAGTCGACTATGCATCCAGCAATCATCAA (f)<br>CCTGCAGGTCGAACTCGAGTTTAGATTGCAGATTCTTCGGCGT (r) | To construct pBad24-CBP-*atcB_Jab*<br>Template: chromosome<br>The PCR product was cloned into XbaI-linearized pBad24-CBPlinker using NEBuilder® HiFi DNA Assembly Cloning Kit. |
| GGGCACTTGTCGAGTCGACTATGAAGTCTATTGCTTTGCCG (f)<br>CCTGCAGGTCGAACTCGAGTTTAGAGGTCTTCAAAAAGGTCAT (r) | To construct pBad24-CBP-*atcB_Dma*<br>Template: chromosome<br>The PCR product was cloned into XbaI-linearized pBad24-CBPlinker using NEBuilder® HiFi DNA Assembly Cloning Kit. |
| GGGCACTTGTCGAGTCGACTATGTCCGACCCAGCCAACCC (f)<br>CCTGCAGGTCGAACTCGAGTTTAGAATCTGTCAAAGACGCCGG (r) | To construct pBad24-CBP-*atcB_Veg*<br>Template: chromosome<br>The PCR product was cloned into XbaI-linearized pBad24-CBPlinker using NEBuilder® HiFi DNA Assembly Cloning Kit. |
| GGGCACTTGTCGAGTCGACTATGATTGTCGCCTTGGCTGA (f)<br>CCTGCAGGTCGAACTCGAGTTTAGAATCTGTCAAAGACGCCGG (r) | To construct pBad24-CBP-*atcB_Tsy*<br>Template: chromosome<br>The PCR product was cloned into XbaI-linearized pBad24-CBPlinker using NEBuilder® HiFi DNA Assembly Cloning Kit. |
| GCCACTGCAGGTCGACTCTAATGAAGGATTATTACGCCGTT (f)<br>GTATCGATAAGCTTGATATCTTATTGCAAAACCTTGTTCATA (r) | To construct pT18-*atcJ_Care*<br>Template: chromosome<br>The PCR product was cloned into EcoRI-linearized pUT18Clinker using NEBuilder® HiFi DNA Assembly Cloning Kit. |

| | |
|---|---|
| GCCACTGCAGGTCGACTCTAATGAAAGCAGCAAACCTGAT (f)<br>GTATCGATAAGCTTGATATCTTACAGGGCGGATTCCTCGCC (r) | To construct pT18-*atcB_Care*<br>Template: chromosome<br>The PCR product was cloned into EcoRI-linearized pUT18Clinker using NEBuilder® HiFi DNA Assembly Cloning Kit. |
| GGGCTGCAGGGTCGACTCTAATGTCCGACAAGAACCGGCAT (f)<br>GTATCGATAAGCTTGATATCTTATTTCTTTTCCTTGCCGGCGT (r) | To construct pT25-*atcC_Care*<br>Template: chromosome<br>The PCR product was cloned into EcoRI-linearized pKT25linker using NEBuilder® HiFi DNA Assembly Cloning Kit. |
| GCCACTGCAGGTCGACTCTAATGCAAGATCATTATGCAAA (f)<br>GTATCGATAAGCTTGATATCTTACTGGATGATTCCTTTAA (r) | To construct pT18-*atcJ_Jab*<br>Template: synthetic gene<br>The PCR product was cloned into EcoRI-linearized pUT18Clinker using NEBuilder® HiFi DNA Assembly Cloning Kit. |
| GCCACTGCAGGTCGACTCTAATGCATCCAGCAATCATCAA (f)<br>GTATCGATAAGCTTGATATCTTAGATTGCAGATTCTTCGG (r) | To construct pT18-*atcB_Jab*<br>Template: synthetic gene<br>The PCR product was cloned into EcoRI-linearized pUT18Clinker using NEBuilder® HiFi DNA Assembly Cloning Kit. |
| GCCACTGCAGGTCGACTCTACTTATCGACAATCCATTAGA (f)<br>GTATCGATAAGCTTGATATCGAGCGGATACATATTTGAATG (r) | To construct pT18-*CteratcJ_Jab*<br>Template: synthetic gene<br>The PCR product was cloned into EcoRI-linearized pUT18Clinker using NEBuilder® HiFi DNA Assembly Cloning Kit. |
| GGGCTGCAGGGTCGACTCTAATGAGCGTAGTGTCTACCAC (f)<br>GTATCGATAAGCTTGATATCTTAGCTGCGTTCTTTCCAGGT (r) | To construct pT25-*atcC_jab*<br>Template: synthetic gene<br>The PCR product was cloned into EcoRI-linearized pKT25linker using NEBuilder® HiFi DNA Assembly Cloning Kit. |
| GCCACTGCAGGTCGACTCTAATGAATTATTACGATATTTTG (f)<br>GTATCGATAAGCTTGATATCTTATTTAAACAGAGTCAAGAA (r) | To construct pT18-*atcJ_Dma*<br>Template: synthetic gene<br>The PCR product was cloned into EcoRI-linearized pUT18Clinker using NEBuilder® HiFi DNA Assembly Cloning Kit. |

| | |
|---|---|
| GCCACTGCAGGTCGACTCTAATGAAGTCTATTGCTTTGCC (f)<br>GTATCGATAAGCTTGATATCTTAGAGGTCTTCAAAAAGGTCTT (r) | To construct pT18-*atcB_Dma*<br>Template: synthetic gene<br>The PCR product was cloned into EcoRI-linearized pUT18Clinker using NEBuilder® HiFi DNA Assembly Cloning Kit. |
| GGGCTGCAGGGTCGACTCTAATGACCTTTTTGAAGACCTCTACCCA (f)<br>GTATCGATAAGCTTGATATCTTACCTACAATTCTTATGAAATCT (r) | To construct pT25-*atcC_Dma*<br>Template: synthetic gene<br>The PCR product was cloned into EcoRI-linearized pKT25linker using NEBuilder® HiFi DNA Assembly Cloning Kit. |
| GCCACTGCAGGTCGACTctaGCTGATGCGGATTCCTTTTTG (f)<br>GTATCGATAAGCTTGATATCGAGCGGATACATATTTGAATG (r) | To construct pT18-Cter*atcJ_Dma*<br>Template: pBad33-*atcC*+ *atcJ_Dma*<br>The PCR product was cloned into EcoRI-linearized pUT18Clinker using NEBuilder® HiFi DNA Assembly Cloning Kit. |
| AACCGAGGGCGCTTACGCTCTGG (f)<br>TCAGCAAGGCTGACCAGTG (r) | To introduce D28A mutation in AtcB_Care |
| CCGCGAATACGCCACCTCGAAAG (f)<br>ATGACGGCCAGCAAATCC (r) | To introduce D51A mutation in AtcB_ Care |
| GTCAGAGTTTCAGCCCGATAGAA (f)<br>GCTTTCTTGCGATAGGC (r) | To introduce H31Q mutation in AtcJ_ Care |
| CGAACAAATCAGGACCACTTATATGAAC (f)<br>GCCGTTTCCAGCGGA (r) | To introduce W83R mutation in AtcJ_ Care |
| CCTGCTGGAATAACCGCTGGAAACG (f)<br>CTCCGGCGCCGGTTT (r) | To insert stop codon (TAA) in the sequence coding for AtcJ_ Care at position N74 (AtcJ_Care$_{\Delta Cter}$) |
| TCAGCACGGTGCTTATGCGCTAG (f)<br>TCTGCAATATCTGATAGCTCTTTTTC (r) | To introduce D33A mutation in AtcB_Jab |
| GCGTGAATACGCCTCATCCAAAG (f)<br>ATAATGGCGAGTAAGTCTTTAG (r) | To introduce D56A mutation in AtcB_Jab |
| CGCCAAATACCAGCCCGACAAAA (f)<br>GCATTTTTACGGTAAGCGGATTTAATTAAATC (r) | To introduce H31Q mutation in AtcB_Jab |
| TGCCCGCGACCGCGCCGAAAAATAC (f)<br>ATCTCTAATGGATTGTCGATAAG (r) | To introduce I82R mutation in AtcJ_Jab |
| CCGCGACATTCGCGAAAAATACATTAAAG (f)<br>GCAATCTCTAATGGATTGTC (r) | To introduce A83R mutation in AtcJ_Jab |
| TGCCGAAAAACGCATTAAAGGAATCATCCAG (f)<br>ATGTCGCGGGCAATC (r) | To introduce Y86R mutation in AtcJ_Jab |

| | |
|---|---|
| CCTTATCGACTAACCATTAGAGATTGC (f)<br>CTGCGTTGCCGGTAG (r) | To insert stop codon (TAA) in the sequence coding for AtcJ_ Jab at position N74 (AtcJ_Dma$_{\Delta Cter}$) |
| TGCGGATTCCTCTTTGTTTGACT (f)<br>TCAGCTACCAATTCCAAAC (r) | To introduce F75S mutation in AtcJ_Dma |
| TTCCTTTTTGTCTGACTACTTCTTG (f)<br>TCCGCATCAGCTACCAAT (r) | To introduce F77S mutation in AtcJ_Dma |
| TTTGTTTGACTCCTTCTTGACTCTGTTTAAATAAG (f)<br>AAGGAATCCGCATCAGCT (r) | To introduce Y79S mutation in AtcJ_Dma |
| GTTTGACTACTCCTTGACTCTGTTTAAATAAG (f)<br>AAAAAGGAATCCGCATCAG (r) | To introduce F80S mutation in AtcJ_Dma |
| CTTGACTCTGTCTAAATAAGATATC (f)<br>AAGTAGTCAAACAAAAAGGAATC (r) | To introduce F84S mutation in AtcJ_Dma |
| GGAATTGGTATAAGATGCGGATTCCTTTTTG (f)<br>AAACGTCCTTGTGAATC (r) | To insert stop codon in (TAA) in the sequence coding for AtcJ_Dma at position A70 (AtcJ_DmaΔCter) |
